# Assessment of Landbird Population Change in the Southeastern United States

**DOI:** 10.1101/2023.05.16.540449

**Authors:** Nicole L. Michel, Dean Demarest, Todd Jones-Farrand, Jeff Gleason, Keith McKnight, Randy Wilson

## Abstract

Birds and their habitats are facing unprecedented threats from a multitude of threats. Threats to birds, and consequently their population trajectories, vary both across space and among species groups, and greater knowledge of these patterns will help inform bird conservation and management. Here, we adapt existing methods to estimate continental bird population loss to a regional scale. Our objectives were to (1) identify patterns in regional population change (abundance) for 141 species of landbirds breeding in the Southeast; (2) compare these with continental patterns for the same suites of species, and (3) examine whether population-level changes in the Southeast suggest immediate, regional conservation actions. We found that landbird population losses were overall similar in the Southeast and across North America (−21%). Shrikes, nightjars, and swifts experienced the largest proportional losses among families in both regions. Birds associated with early seral and emergent wetland habitats experienced the greatest losses as did partial migrants and species listed on the Birds of Conservation Concern list for southeastern Bird Conservation Regions. Facultative aerial insectivores experienced the greatest losses in both regions, while obligate aerial insectivores increased in the Southeast in contrast to continental declines, due to rapid population growth in Cliff Swallows (*Petrochelidon pyrrhonota*). Within the Southeast, the greatest bird losses were in Peninsular Florida and Gulf Coastal Prairie, while the Mississippi Alluvial Valley Bird Conservation Region – where extensive reforestation efforts have been undertaken – had the smallest losses. We found clear differences in patterns of landbird population loss between the Southeastern United States and North America, as well as within the Southeast region. Results from these analyses should provide conservation agencies and partnerships with additional information and new perspectives to guide landscape-level planning and on-the-ground bird conservation delivery efforts to help bring back the nearly three billion birds lost across North America since 1970.

**Disclaimer:** The findings and conclusions in this article are those of the authors and do not necessarily represent the views of the U.S. Fish and Wildlife Service.

## INTRODUCTION

Birds and their habitats are facing unprecedented threats from climate change, habitat degradation and conversion, incompatible land use practices, environmental contamination, collision-related mortality, and non-native predators just to name a few. These threats result directly in mortality, as well as having indirect effects to survival and fitness via complex interactions affecting behavior, physiology, and the ability to acquire resources in completing key stages of the annual cycle (e.g., nesting, molt, migration). The conservation community has worked diligently to conserve birds by mitigating these threats, and their efforts are beginning to payoff for waterfowl which have rebounded thanks to coordinated, landscape level wetland conservation efforts (North American Bird Conservation Initiative 2022). Yet most North American breeding bird populations have continued to decline, with an estimated 2.9 billion less breeding individuals than in 1970 (Rosenberg et al. 2019). Moreover, many species are projected to lose an additional 50% or more of their present population over the next 50 years (North American Bird Conservation Initiative 2022).

Over that same interval, three taxonomic groups of North American birds (waterbirds, dabbling and diving ducks, and geese and swans) have seen population increases of 18 – >1,000% since 1970 (North American Bird Conservation Initiative 2022). Concurrently, sea ducks, shorebirds, and landbirds have declined by 5% to 34%, and a subset of 70 Tipping Point species – species that have already lost ≥ 50% of their populations and are predicted to lose at least an additional 50% of their current populations in the next 50 years – have declined by over two-thirds (North American Bird Conservation Initiative 2022). Thus, patterns of population change within the North American avifauna are variable across species sharing similar habitats, and within and among taxonomic groups.

Threats to bird populations also vary regionally and may differentially contribute to changes in the population status of species continentally. It can be informative and therefore important to elucidate and address these regional variations. For example, the total biomass of spring northerly migrant birds declined in the Mississippi and Atlantic flyways during 2007-2017, while no change was observed in flyways further west (Rosenberg et al. 2019). This pattern may represent threats such as habitat loss and fragmentation on the breeding and/or wintering grounds differentially affecting the temperate- and boreal-breeding landbirds that dominate migrants in these more eastern flyways. Even within the Mississippi and Atlantic Flyways, long-term (90-year) change in winter bird occurrence probabilities varied both spatially and among nine habitat-based species groups (Saunders et al. 2022).

Bird populations may also be differentially affected by threats operating during distinct phases of their annual cycle. For example, Connecticut Warblers (*Oporornis agilis*) and Wood Thrushes (*Hylocichla mustelina*) are predominantly impacted by breeding habitat loss (Rushing et al. 2016, Hallworth et al. 2021), while Golden-winged Warbler (*Vermivora chrysoptera*) population trends have been linked to habitat loss on the wintering grounds (Kramer et al. 2018). Because threats may differentially influence bird populations across space and time, it is important to understand potential patterns at regional or less than range-wide scales as a means of exploring drivers of range-wide population change. Examining regional-level patterns in bird population change can be important in elucidating and understanding threats underpinning range-wide population changes thereby providing insight to guide conservation actions.

A robust understanding of region-specific bird population trajectories will help strengthen the scientific basis of conservation and management efforts undertaken by a host of conservation partners. Under the Migratory Bird Treaty Act, the United States Fish and Wildlife Service (USFWS) has statutory authority (16 U.S. C. 703-712;) for the conservation and management of migratory birds in the United States and territories. As such, USFWS relies on baseline regional status and trend information on bird populations, as well as the ability to obtain regular updates to evaluate the impacts of their actions and adaptively inform conservation efforts over time (see Williams et al. 2009, Williams and Brown 2012). Rosenberg et al. (2019) provides a robust method for quantifying absolute changes in the size of breeding bird populations. Moreover, the modular nature of this method permits it to be readily applied to any suite of bird species that are well-surveyed at any scale by standardized annual monitoring programs such as the North American Breeding Bird Survey (Sauer et al. 2020). While Rosenberg et al. (2019) produced compelling estimates summarizing trends and losses in bird populations at the continental scale, regional patterns in bird population trends or the magnitude of gain/loss were not examined.

To better inform regional bird conservation activities in the southeastern United States, (hereafter Southeast), the USFWS sought a transparent and repeatable process for assessing changes in population size (i.e., abundance) in the native avifauna of this region. As a case study, we developed a process for estimating changes in breeding populations of landbirds across the Southeast following the methods published by Rosenberg et al. (2019). Our objectives were to: (1) identify patterns in regional population change (abundance) for landbirds breeding in the Southeast; (2) compare these with continental patterns for the same suites of species, and (3) examine whether population-level changes in the Southeast suggest immediate, regional conservation actions. More specifically, we tested the null hypothesis that overall patterns of population change for breeding landbirds in the Southeast would not differ from continental estimates derived for the same species groups using the approach of Rosenberg et al. (2019). Gaining an improved understanding of regional population change in the Southeast (gains or losses) should aid in more clearly elucidating species, guilds, or communities of birds whose breeding populations may be responding differentially at regional and continental levels and may warrant some form of unique regional attention.

## METHODS

### Study Area

We assessed changes in breeding bird populations across eight Bird Conservation Regions (BCR; Fig. 1): Central Hardwoods (BCR 24), West Gulf Coastal Plain / Ouachitas (BCR 25), Mississippi Alluvial Valley (BCR 26), Southeastern Coastal Plain (BCR 27), Appalachian Mountains (BCR 28), Piedmont (BCR 29), Peninsular Florida (BCR 31), and Gulf Coastal Prairie (BCR 37) that overlap the USFWS Southeast Region administrative boundary. BCRs define regions with similar habitat types, climatic conditions, and bird communities (Bird Studies Canada and North American Bird Conservation Initiative 2014), making them suitable population units for conservation planning and analysis.

**Fig. 1.**
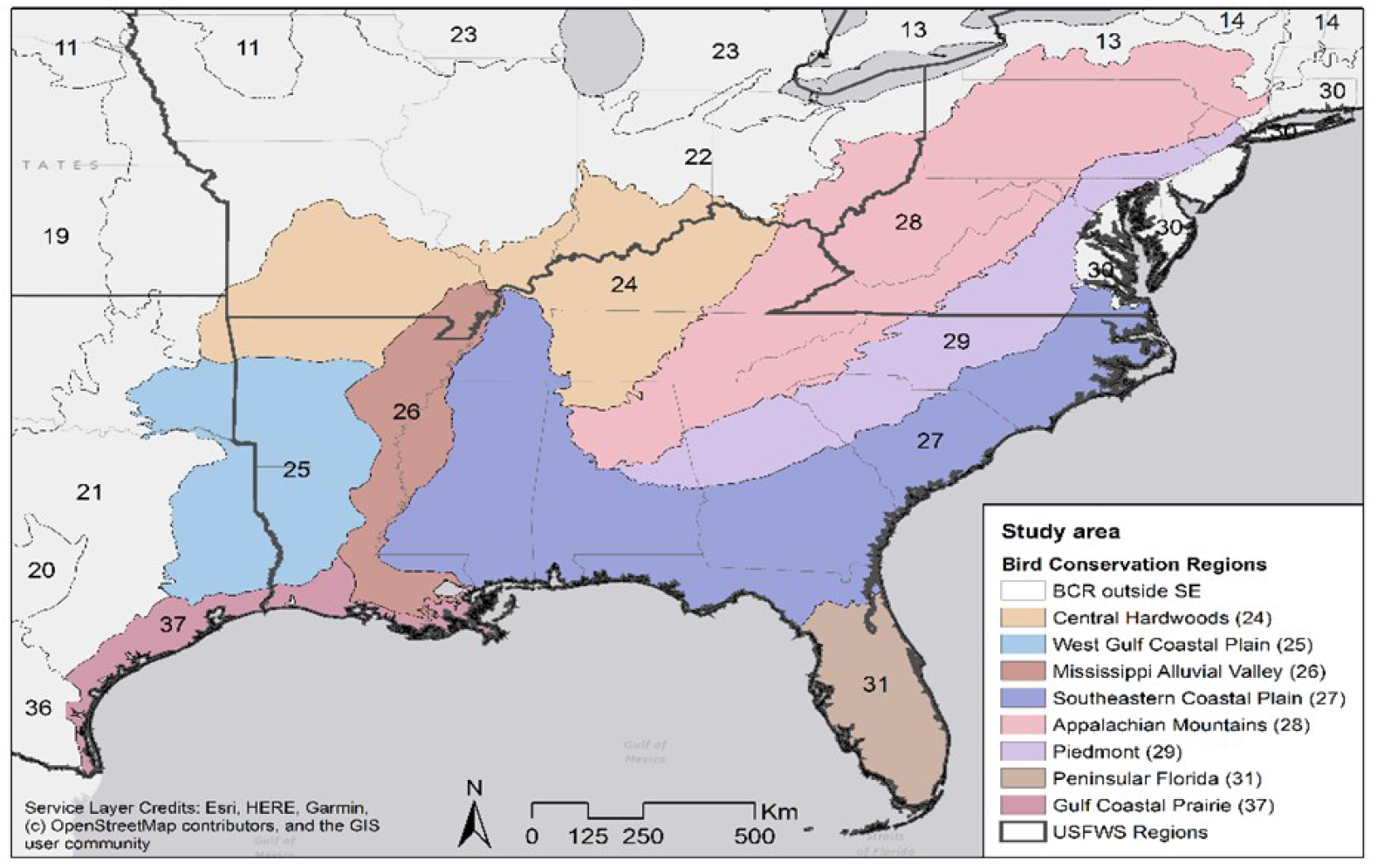
Study area map showing the eight Bird Conservation Regions that intersect with the U.S. Fish and Wildlife Service Southeastern Region.

### Data

Like Rosenberg et al. (2019), our analysis employed two principal data sets: (1) North American Breeding Bird Survey annual abundance indices from 1970-2019 (Sauer et al. 2020), and population estimates from the Partners in Flight (PIF) Population Estimates Database (Partners in Flight 2020, Will et al. 2020). Unlike Rosenberg et al. (2019), we did not employ additional datasets (e.g., eBird) to estimate trends for species where BBS data were unavailable or insufficient (see Study Species). We set 1970 as our starting year following Rosenberg et al. (2019). We used annual indices of abundance derived from Bayesian hierarchical trend models (Sauer and Link 2011) applied to data collected in a “Core” survey area (i.e., from strata [defined for BBS trend models as unique combinations of BCRs and states or provinces] in which one or more routes had data collection extending from 1968 to 2019). These indices provide a full population trajectory showing change in estimated route-level abundance index over the 50-year time series, and thus provide more information than numeric trends (i.e., percent increase or decrease) alone. PIF population estimates were derived using BBS survey data from 2006-2015 and species-specific correction factors following the methods described by Stanton et al. (2019). We used annual indices and population estimates summarized at the BCR level, the smallest spatial scale for which derived BBS annual indices were made publicly available. While BBS data are analyzed at a smaller scale (strata = unique combinations of BCRs and states), we chose not to use summaries at this scale as they have more limited data and consequently, large associated error estimates, making inference challenging. We incorporated estimates of uncertainty from both BBS annual indices and PIF population estimates.

### Study Species

Species considered for analysis included 185 landbirds for which both BBS annual indices of abundance and PIF population estimates were available for one or more of the eight BCRs overlapping the Southeast. Each species was reviewed to ensure: (1) robustness of the annual indices of abundance based upon the number of detections within a BCR (i.e., n ≥ 25 detections), and (2) species breeding within the BCR occurred in sufficient abundance to be considered a “manageable population” in the BCR. Landbird species excluded from analysis on the basis of insufficient detections typically represented species considered to be vagrants to the Southeast, or species occurring along range boundaries (e.g., Hooded Oriole [*Icterus cucullatus*], Verdin [*Auriparus flaviceps*], Vermillion Flycatcher [*Pyrocephalus obscurus*], etc.). Additionally, we excluded species that breed in manageable numbers in BCRs overlapping the USFWS Southeast Region, but whose occurrence in those BCRs is outside of or not representative of the USFWS Southeast Region (e.g., Mourning Warbler [*Geothlypis philadelphia*], Bobolink [*Dolichonyx oryzivorus*]). Species that are largely indistinguishable in the field (e.g., Alder Flycatcher [*Empidonax alnorum*] and Willow Flycatcher [*E. traillii*]) were lumped in the BBS data, per Sauer et al. (2020) and Rosenberg et al (2019) and subsequently treated as a single species for these analyses. Based on the above criteria, we retained 146 of original 185 species evaluated; these 146 species represented 36 families (Appendix 1).

Preliminary analysis demonstrated that Blue-gray Gnatcatcher (*Polioptila caerulea*) accounted for >16% (i.e., 168 million of the 1,016 billion estimated individuals) of the total summed population of the 146 species in our study (Appendix 1, Fig. 2). Because we used a hierarchical modeling approach that draws power from combining population trajectories from many species, we were concerned that Blue-gray Gnatcatcher could disproportionately influence group-level estimates of population change and trajectories due to their disproportionate abundance in the study area (see Fig. 2). Consequently, we retained this species only for the all-species regional population change summary to facilitate comparison with continental-level change estimates, as this species was included in the Rosenberg et al. (2019) continental analysis. Subsequently, we removed Blue-gray Gnatcatcher from all group-level and BCR analyses (see below) to eliminate the potential influence of this single, highly abundant species on inferences regarding group-level patterns of population change within the Southeast.

**Fig. 2.**
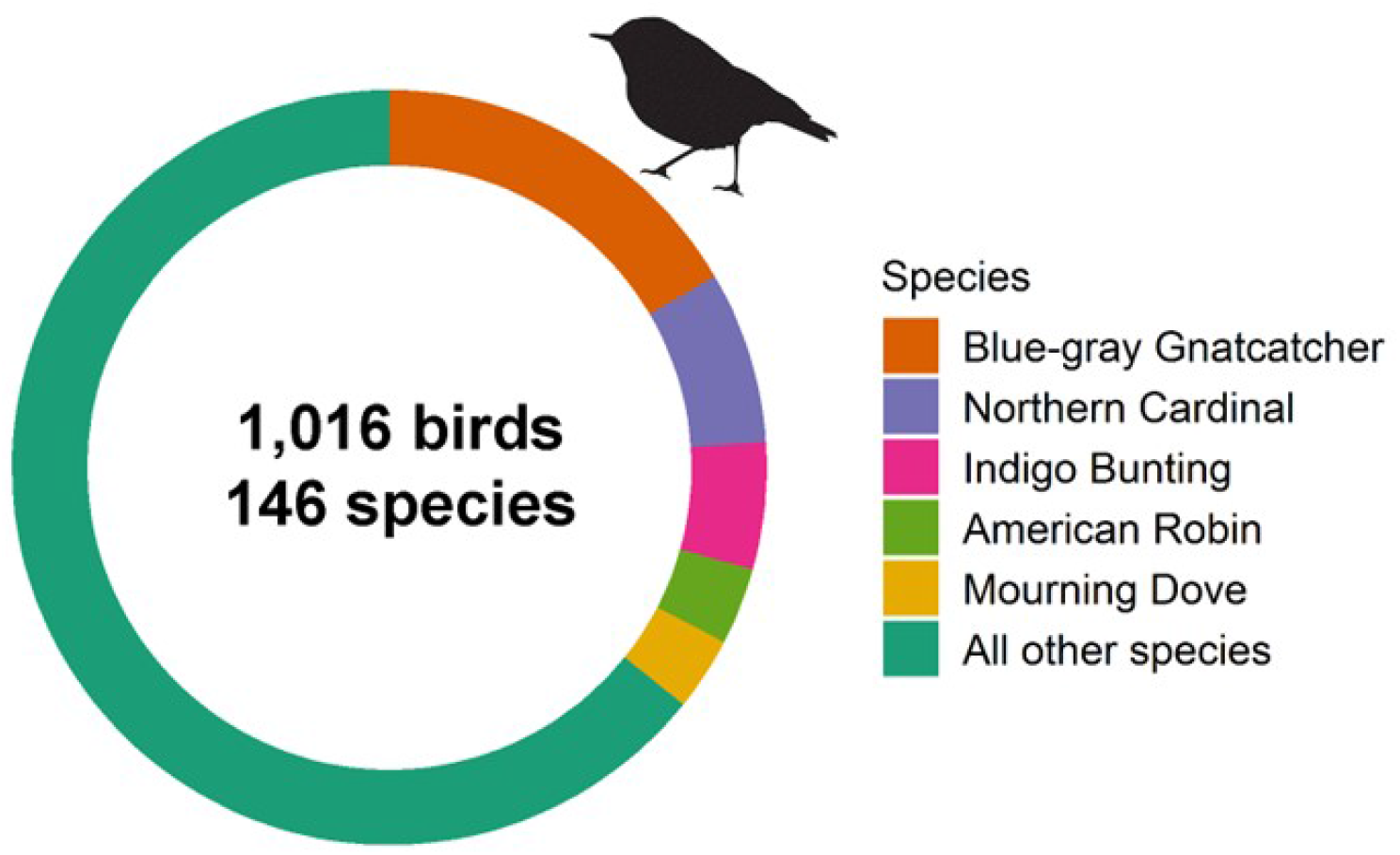
Proportion of summed population estimate for all landbirds in the Southeastern US (n = 146 species) explained by the five most abundant species and all other species combined. Blue-gray Gnatcatcher accounts for ∼16% of the total Southeastern landbird population (i.e., 168 of 1,016 million estimated individuals).

Similarly, our preliminary analyses included four highly abundant introduced species that have become established in the Southeast: Eurasian Collared-Dove (*Streptopelia decaocto*), European Starling (*Sturnus vulgaris*), House Sparrow (*Passer domesticus*), and Rock Pigeon (*Columba livia*). Because these non-native species do not fall under the purview of USFWS as migratory birds in the public trust but were included in the Rosenberg et al. (2019) analysis, we retained them only in the all-species summary analysis at the regional level (i.e., for comparison with the continental estimates), but excluded them from group-level and BCR analyses within the Southeast. Thus, after excluding introduced species and Blue-gray Gnatcatcher, our final group-level and BCR analyses within the Southeast incorporated 141 landbird species representing 33 families.

### Species Groupings

For purposes of examining patterns of population change across the Southeast, all species (n = 141) were assigned to five groups for further analyses: (1) taxonomic family; (2) habitat association; (3) migratory status; (4) conservation status; and (5) aerial forager status (Appendix 2: Table A2.1). Because each species was assigned to each of the five analysis groups (Appendix 1), all group-level analyses are independent of each other. Group assignments for habitat association, migratory status, and aerial foraging were based on published literature and USFWS subject matter expert recommendations.

- *Taxonomic Family*: All 141 species were categorized according to their taxonomic family (AOS 2021), resulting in 33 mutually exclusive, family-level categories.
- *Habitat Associations*: Habitat associations consisted of two hierarchical classes (Level I and Level II). Level I associations represent the general landcover type(s) individual species were associated with across the Southeast (e.g., aerial, emergent wetland, forest woodland, generalist, grassland/open land, and shrub-scrub & successional). Whereas Level II associations represent finer-scale landcover types nested within Level I habitat associations (e.g., Level I = forest woodland and Level II = eastern deciduous, southern pine, and spruce-fir/northern hardwood and forest woodland generalist). Bird species are often associated with multiple Level I and Level II habitat associations and consequently, their regional pattern of population change contributes to the overall pattern for multiple habitat associations. As such, overall population change for a given habitat association indicates change for a suite of birds associated with that habitat; some of which may also be contributing to patterns of change in other habitat associations. That is, the pattern of change for a given species across the entire Southeast may influence the overall pattern for a given habitat even though the species may not occur exclusively in that habitat.
- *Migratory Status*: All species were assigned to one of three mutually exclusive categories of migratory status: migrant, partial migrant, and resident. Migrants were defined as species whose breeding and wintering ranges do not overlap, partial migrants as species that migrate but with overlapping breeding and wintering ranges, and residents as species who remain largely within a defined geography year-round.
- *Conservation Status*: We evaluated species based on a coarse assessment of conservation vulnerability that grouped species into two mutually exclusive categories, Birds of Conservation Concern (BCC) and non-BCC species. BCC species were those identified on the most recent BCC report (U.S. Fish and Wildlife Service 2021) as either species of continental concern, species of concern for one or more BCRs in the Southeast, or species of concern for one or more BCRs outside the Southeast.
- *Aerial Insectivores*: Given recent attention to demonstrated declines among aerial insectivores (e.g., Nebel et al. 2010; Spiller and Dettmers 2019) we also grouped species into three categories based on aerial foraging behavior. Species were categorized as obligate aerial foragers (species that exclusively forage in air space, [e.g., Chimney Swift *Chaetura pelagica*)], facultative aerial foragers (species that regularly forage in air space as well as other microhabitats, [e.g., Acadian Flycatcher Empidonax virescens]), or non-aerial foragers (species whose primary mode of foraging does not involve aerial habitats; [e.g., American Robin *Turdus migratorius*]).

### Population Change Estimation

We adapted the analytical methods and R code published in Rosenberg et al. (2019), which produced population change estimates for all of North America. We produced population change estimates for 141 – 146 landbird species (see Study Species section) and five custom groups (see Species Groupings) across the Southeast region, as well as for each of eight BCRs overlapping the region. All analyses were conducted in R version 4.0.5 (R Core Team 2021).

We extracted BBS annual indices of abundance (1970-2019) summarized at the BCR level (Sauer et al. 2020) for the eight BCRs in the Southeast (Fig. 1). We then conducted a seven-step modeling procedure that downscaled the methods from Rosenberg et al. (2019) to the Southeast. First, for each species individually, we applied a hierarchical Bayesian generalized additive model (GAM) to smooth the population trajectory. This step was done as a precursor to the all-species hierarchical model because species exhibit a high degree of inter-annual variation in their population trajectories. This smoothing step maintained each species’ overall trajectory while reducing the magnitude of population fluctuations, so that the all-species model was not unduly influenced by species with highly variable population trajectories. We used a highly flexible GAM (k = 11 knots) implemented in mgcv (Wood 2011) that incorporated the estimates of uncertainty (i.e., 95% confidence intervals) that accompanied the BBS annual indices. Because we produced total and group-level population change estimates for each BCR separately (see step five), we conducted this smoothing step at the scale of the BCR. That is, separate models were run for each species for each BCR it occurred in (i.e., up to 8 GAMs per species). We applied the GAM smoother to natural log-transformed BCR-level trajectories separately to ensure that the same smoothed population trajectories were used in the estimates of population change for both the Southeast and for each BCR individually.

Second, we produced a single regional population trajectory per species by summing across all the BCRs for which the species is known to occur. We summed the smoothed annual indices produced by the single-species BCR-level GAMS (from step one) for each species, and standardized the indices relative to 1970 (i.e., set 1970 to equal 1 and adjusted the other indices accordingly). We calculated summed standard errors for regional trajectories as the square root of the summed variances from each BCR-level trajectory and converted them to 95% confidence intervals.

Third, we fit a single, all-species hierarchical Bayesian model to estimate annual population size (and consequently gain or loss) for each species and species group while accounting for influences across the full annual cycle (e.g., Hostetler et al. 2015). Because our objective was prediction rather than hypothesis testing, we followed Rosenberg et al. (2019) in fitting a single full model rather than a suite of alternate models. To accomplish this, we incorporated influences across the full annual cycle by combining information among species that shared each unique combination of breeding and non-breeding biomes (see Appendix 1), based on the assumption that species using the same regions and habitats would face similar pressures. Breeding and non-breeding biomes were not otherwise treated as groupings for assessment as described above. The smoothed, standardized regional trajectories for each species (from steps one and two, above) comprised the data entering this single, all-species hierarchical model. Species-specific smoothed regional trajectories were log-transformed to put each species on a common scale and modeled as normal random variables with variances drawn from their 95% confidence intervals. Means were determined by a hyperparameter representing mean status for each breeding/non-breeding biome group with an associated variance. Mean trajectories for each breeding/non-breeding biome group were themselves estimated using an additive submodel that used random walk time-series to model the effects of each seasonal biome. Random walks were used as they allow non-linear change while helping smooth large interannual fluctuations (Rosenberg et al. 2019).

This all-species hierarchical model also incorporated each species’ regional population size estimate. We produced a single regional population estimate per species by summing BCR-level population estimates and their associated error for 2006-2015 (Partners in Flight 2020). Selection of the 2006-2015 time period allows for both backward and forward projection of population estimates while incorporating associated 95% CIs around the annual indices and the population estimate, respectively. Population size was incorporated as random draws from a normal distribution based on the published estimates (Partners in Flight 2020). We used the log-transformed standardized population trajectories from 2006-2015 to calculate a scaling factor that we applied to the entire population trajectory to generate annual population size estimates. The model was implemented in Jags using R packages *rjags* (Plummer 2019) and *jagsUI* (Kellner 2021) and plotted using *ggplot2* (Wickham 2016).

Fourth, we calculated annual estimates of total (i.e., numerical) population change by subtracting each year’s total population estimate for all species from the corresponding total 1970 population estimate. Proportional (i.e., percentage) changes were also calculated

Fifth, we repeated the previous four steps for each of the five groupings described above (*Species Groupings*). This produced annual estimates of total and proportional population loss between 1970 and 2019 for all birds assigned to a given category within each grouping (e.g., migratory status, conservation status, habitat association) across the Southeast.

Sixth, we repeated the above five-step analytical procedure for each of the eight BCRs individually. To produce BCR-specific population change estimates (i.e., versus estimates for the Southeast overall), we skipped step two, instead retaining the smoothed population trajectory from each BCR rather than summing across the Southeast. We then standardized and log-transformed each BCR’s smoothed population trajectory (step three), and fit separate all-species hierarchical Bayesian models for each BCR. This produced annual estimates of total and proportion population loss/gain for the set of species (*n* = 81 – 129) that occur in each of the respective BCRs.

Finally, to directly compare our Southeast population change estimates with continental estimates, we produced annual estimates of total and proportional population loss/gain for each of the 146 study species at the continental scale. We extracted BBS annual indices of abundance for the same time period (1970-2019) from the same model as above (Sauer et al. 2020) but summarized at the continental scale rather than by BCR. We repeated the first five steps above to produce population change estimates for all species combined, categories within each of the five species groupings, and each species individually.

### Numeric vs. Proportional Comparisons

Due to disparities in spatial scale, we emphasized proportional population change (% population loss/gain since 1970) rather than numerical population change (numbers of individuals) for certain comparisons. For example, changes in numerical abundance for landbird species analyzed were expected to be far greater across North America than in the Southeast, and greater in BCRs of significantly larger spatial extent. Similarly, in comparisons of population change among taxonomic families, or among other categories of species’ groupings (e.g., BCC species versus non-BCC species) where the number of species comprising each category could substantially differ, proportional comparisons were favored as potentially much more revealing of noteworthy patterns and/or possible underlying drivers of population change. Summaries of numeric change for all species and species groupings are presented in Appendix 1 and 3. For all comparisons, we deemed them different (*P* < 0.05) where respective 95% CIs did not overlap zero (Cherry 1998).

## RESULTS

### Overall Landbird Population Change in the Southeast

Across the Southeast, total landbird abundance declined by an estimated -260.6 million birds (hereafter 95% CIs reported for all medians: -282.6, -238.2; *n* = 146 species) from the approximately one billion breeding bird population in 1970 (Table 1, Fig. 3A). This represents a loss of over one-fifth (−22% [-0.23, -0.20]) of the total breeding landbird population in our study area since 1970, with 48% of the species exhibiting declines.

**Table 1.** Population change among landbird species in the Southeastern US and the same species set across North America, 1970-2019. Number of species, abundance change (in millions), proportion change, and proportion of declining species shown for three composite groups of landbird species for each region. The composite groups represent: all species (n = 146), all species excluding Blue-gray Gnatcatcher (BGGN; n = 145), and all native species excluding BGGN (n = 141).

| Species Group | Number of Species | Southeastern US |  |  |  |  | North America |  |  |  |  |
| --- | --- | --- | --- | --- | --- | --- | --- | --- | --- | --- | --- |
|  |  | Numerical Abundance Change (Millions) |  | Proportion Abundance Change |  | Proportion Declining Species | Numerical Abundance Change (Millions) |  | Proportion Abundance Change |  | Proportion Declining Species |
|  |  | Change | 95% CI | Change | 95% CI |  | Change | 95% CI | Change | 95% CI |  |
| All species | 146 | -260.63 | (-282.66, -238.20) | -0.22 | (-0.23, -0.20) | 0.48 | -1331.78 | (-1422.89, -1237.33) | -0.24 | (-0.25, -0.22) | 0.57 |
| Excluding BGGN | 145 | -270.69 | (-288.38, -252.26) | -0.25 | (-0.27, -0.24) | 0.48 | -1368.21 | (-1460.12, -1275.11) | -0.25 | (-0.27, -0.24) | 0.57 |
| Excluding BGGN + non-natives | 141 | -199.07 | (-215.10, -182.74) | -0.21 | (-0.22, -0.19) | 0.50 | -1041.50 | (-1121.40, -962.47) | -0.21 | (-0.23, -0.20) | 0.57 |

**Fig. 3.**
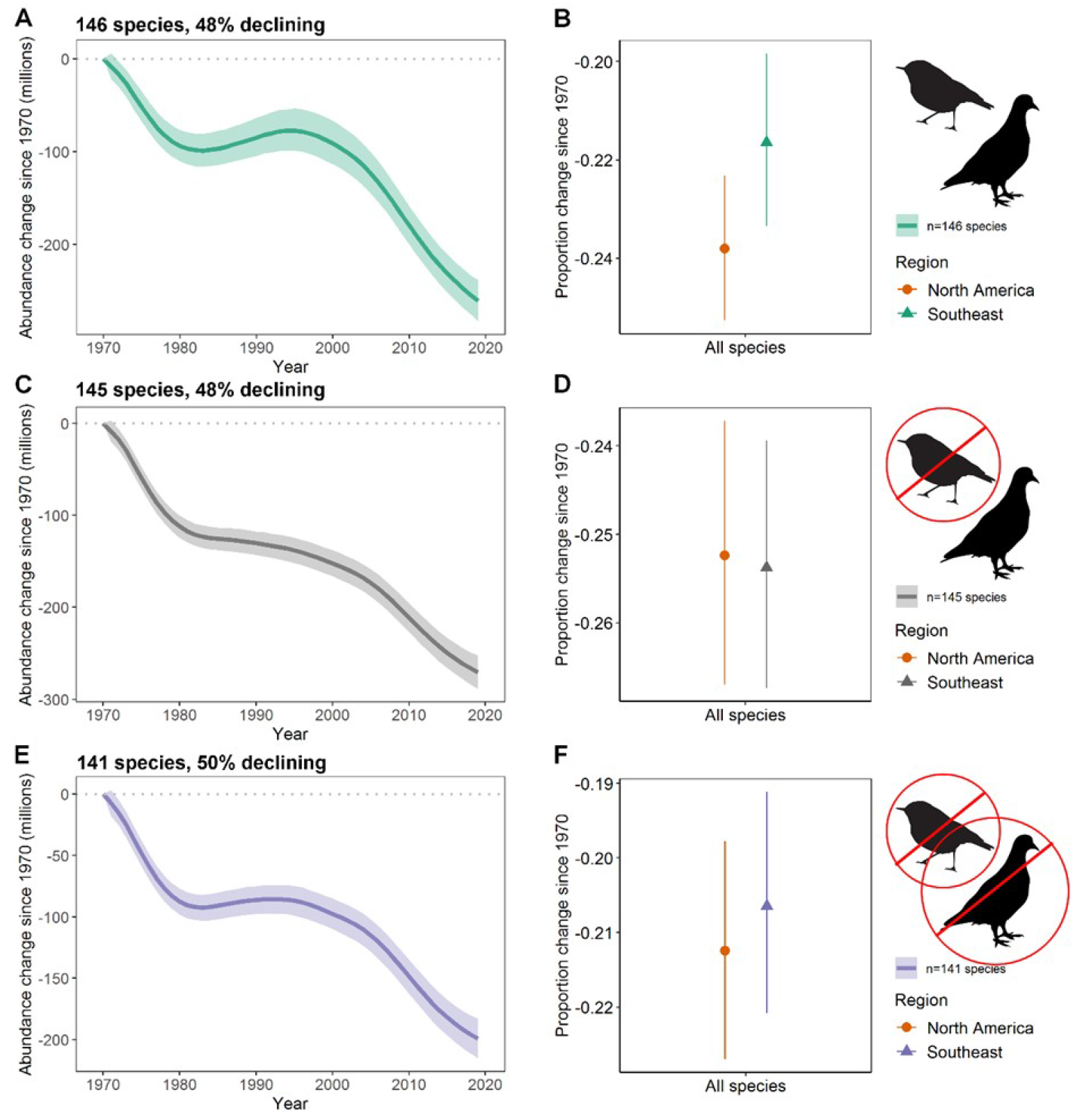
Net change in abundance of landbirds across the Southeastern US during 1970-2019 (A, C, E), and comparisons with proportional abundance change across North America during the same time period (B, D, F) for different suites of species. Change estimates are shown for all landbirds that met our criteria for inclusion (see Methods; n = 146 species; A, B); all landbirds excluding Blue-gray Gnatcatcher (n = 145 species; C, D); and all landbirds excluding Blue–gray Gnatcatcher and introduced species (n = 141 species; E, F). Annual median population change estimates (in millions) (A, C, E) or estimates of proportional change between

After removing Blue-gray Gnatcatcher from the analysis, total breeding landbird abundance for the remaining 145 species declined by -270.69 million (−288.38, -252.26), a loss of one-quarter (−25% [-0.27, -0.24]) of the landbird population with 50% of the species exhibiting population declines (Table 1, Fig. 3C). The population trajectory (Fig. 3C) showed a more linear decline with less of an increase during the 1980s and 1990s compared to that for all species (Fig. 3A), suggesting that the pattern for all 146 species was strongly influenced by an increasing trend in the very abundant Blue-gray Gnatcatcher during these decades.

After removing Blue-gray Gnatcatcher and all non-native bird species from the analysis, total breeding landbird abundance for the remaining 141 native bird species declined by -199.07 million individuals (−215.10, -182.74; Table 1). This represents a loss of over one-fifth of the native landbird population (−21% [-0.22, -0.19]), with 50% of native bird species declining (Fig. 3E). The population trajectory for native landbirds alone exhibited an increase in the 1980s (Fig. 3E) relative to the trajectory for the suite of 145 species that only excluded Blue-gray Gnatcatcher (Fig. 3C), suggesting that declines in the four abundant introduced bird species strongly influenced the pattern during the 1980s when included in the analysis (i.e., ameliorated an otherwise slightly increasing pattern in native bird species during this decade).

### Overall Landbird Population Change in the Southeast Versus North America Estimates

The Southeast lost a slightly smaller proportion of the total population of 146 landbird species than were lost across North America (−0.24% [-0.25, -0.22]; Fig. 3B). After removing Blue-gray Gnatcatcher from the comparison, the Southeast and North America (−25% [-0.27, -0.24]) exhibited similar proportional losses across 145 species (Fig. 3D). On removing non-native species in addition to Blue-gray Gnatcatcher, proportional landbird population losses for the resultant 141 species remained similar for the Southeast and North America (−21% [-0.23, -0.20]; Fig. 3F).

### Landbird Population Change by Species Groupings

#### Southeast

The largest numerical change in abundance among landbird families in the Southeast was within the Icteridae (New World Blackbirds; *n* = 9 species), with losses of -119.3 million birds ([-130.1, -109.0]; Table 2, Fig. 4B). Over 60% of the losses in Icteridae were attributable to declines in Eastern Meadowlarks (*Sturnella magna*), which lost -39.7 million birds (−46.3, -33.3), and Common Grackles (*Quiscalus quiscula*), which lost -38.9 million birds (−44.8, -33.1; Appendix 1). Among non-Passerines in the Southeast, Caprimulgidae (Nightjars; n = 3 species) exhibited the greatest change in total abundance, with a loss of -15.3 million birds ([-18.0, -12.7]; Table 2, Fig. 4A). All three species of nightjars included in this study declined since 1970, with Chuck-will’s-widow (*Antrostomus carolinensis*) comprising the greatest losses at -7.0 million birds ([-8.3, 5.7]; Appendix 1). Conversely, the greatest numeric population gains in the Southeast occurred within Hirundinidae (Swallows and Martins; n = 6 species) with a gain of 14.8 million birds (9.1, 23.0). However, 85% of this increase was attributed to a gain of 12.7 million birds of a single species, Cliff Swallow (*Petrochelidon pyrrhonota* [7.2, 20.4]; Appendix 1). Large population gains were also observed in Vireonidae (Vireos; *n* = 6 species) with a gain of 14.7 million birds ([12.2, 17.3]; Table 2).

**Table 2.**
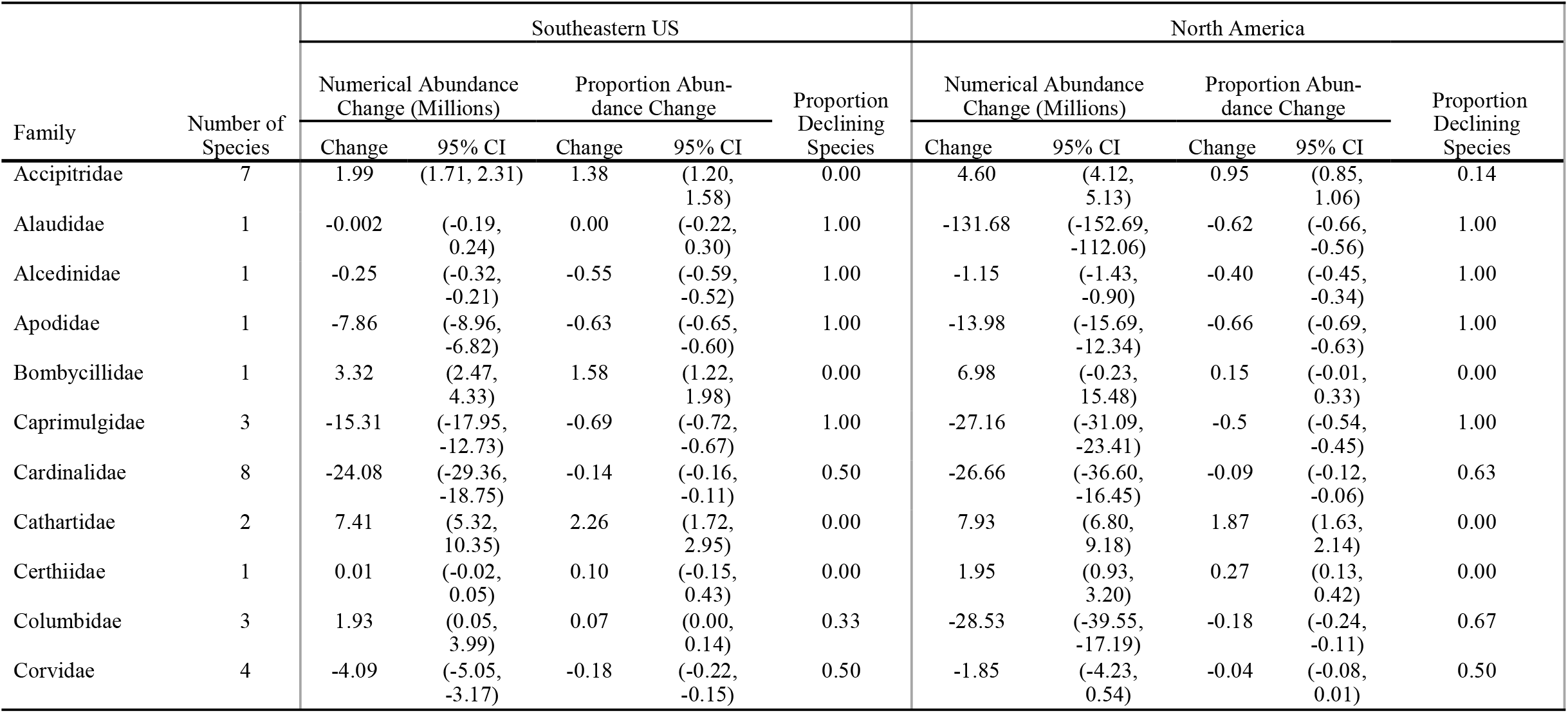
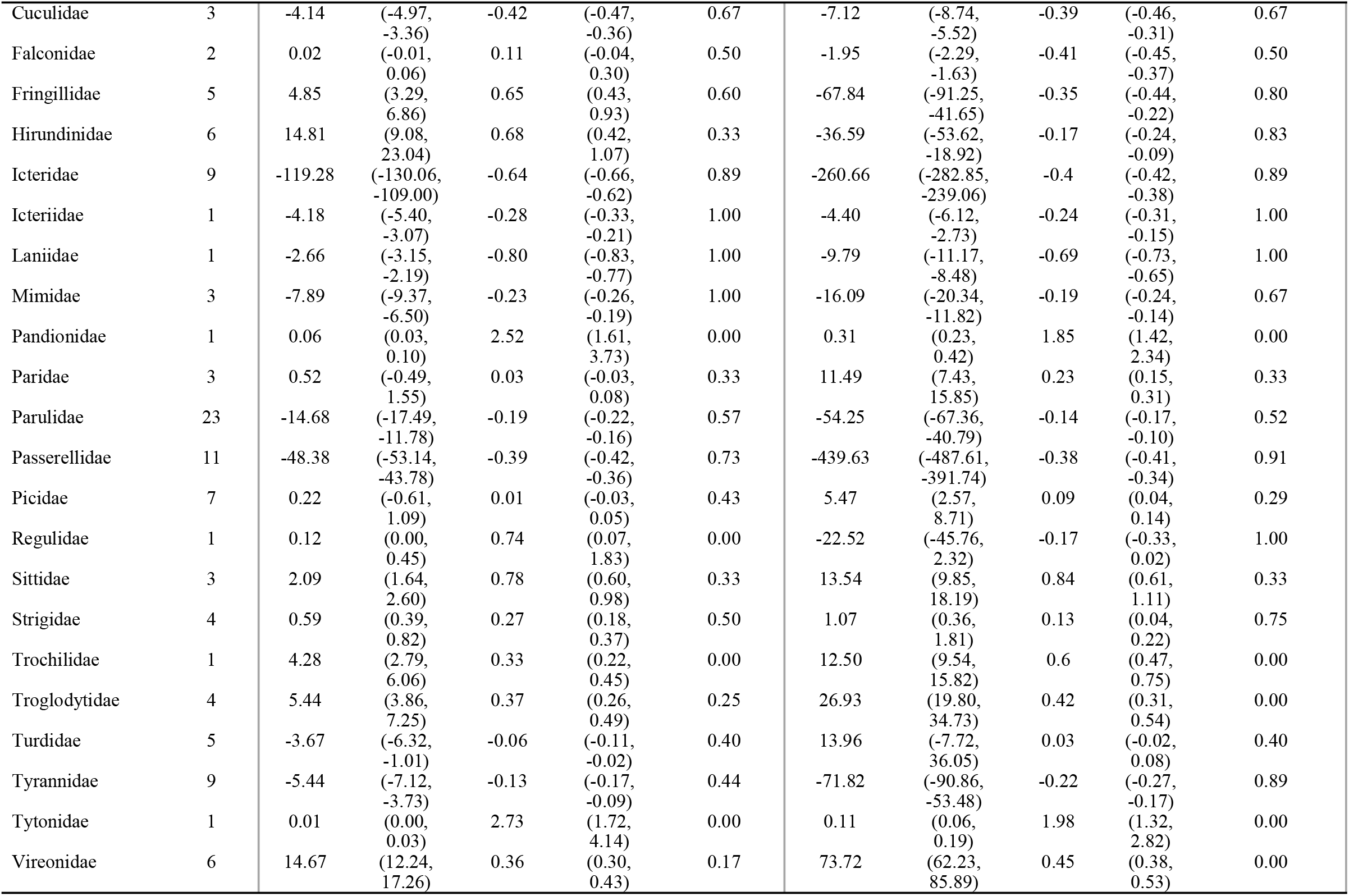
Population change by taxonomic family of landbirds in the Southeastern US and the same species set across North America, 1970-2019. Number of species, abundance change (in millions), proportion change, and proportion of declining species shown for all native species excluding BGGN (n = 141).

| Family | Number of Species | Southeastern US |  |  |  |  | North America |  |  |  |  |
| --- | --- | --- | --- | --- | --- | --- | --- | --- | --- | --- | --- |
|  |  | Numerical Abundance Change (Millions) |  | Proportion Abundance Change |  | Proportion Declining Species | Numerical Abundance Change (Millions) |  | Proportion Abundance Change |  | Proportion Declining Species |
|  |  | Change | 95% CI | Change | 95% CI |  | Change | 95% CI | Change | 95% CI |  |
| Accipitridae | 7 | 1.99 | (1.71, 2.31) | 1.38 | (1.20, 1.58) | 0.00 | 4.60 | (4.12, 5.13) | 0.95 | (0.85, 1.06) | 0.14 |
| Alaudidae | 1 | -0.002 | (-0.19, 0.24) | 0.00 | (-0.22, 0.30) | 1.00 | -131.68 | (-152.69, -112.06) | -0.62 | (-0.66, -0.56) | 1.00 |
| Alcedinidae | 1 | -0.25 | (-0.32, -0.21) | -0.55 | (-0.59, -0.52) | 1.00 | -1.15 | (-1.43, -0.90) | -0.40 | (-0.45, -0.34) | 1.00 |
| Apodidae | 1 | -7.86 | (-8.96, -6.82) | -0.63 | (-0.65, -0.60) | 1.00 | -13.98 | (-15.69, -12.34) | -0.66 | (-0.69, -0.63) | 1.00 |
| Bombycillidae | 1 | 3.32 | (2.47, 4.33) | 1.58 | (1.22, 1.98) | 0.00 | 6.98 | (-0.23, 15.48) | 0.15 | (-0.01, 0.33) | 0.00 |
| Caprimulgidae | 3 | -15.31 | (-17.95, -12.73) | -0.69 | (-0.72, -0.67) | 1.00 | -27.16 | (-31.09, -23.41) | -0.5 | (-0.54, -0.45) | 1.00 |
| Cardinalidae | 8 | -24.08 | (-29.36, -18.75) | -0.14 | (-0.16, -0.11) | 0.50 | -26.66 | (-36.60, -16.45) | -0.09 | (-0.12, -0.06) | 0.63 |
| Cathartidae | 2 | 7.41 | (5.32, 10.35) | 2.26 | (1.72, 2.95) | 0.00 | 7.93 | (6.80, 9.18) | 1.87 | (1.63, 2.14) | 0.00 |
| Certhiidae | 1 | 0.01 | (-0.02, 0.05) | 0.10 | (-0.15, 0.43) | 0.00 | 1.95 | (0.93, 3.20) | 0.27 | (0.13, 0.42) | 0.00 |
| Columbidae | 3 | 1.93 | (0.05, 3.99) | 0.07 | (0.00, 0.14) | 0.33 | -28.53 | (-39.55, -17.19) | -0.18 | (-0.24, -0.11) | 0.67 |
| Corvidae | 4 | -4.09 | (-5.05, -3.17) | -0.18 | (-0.22, -0.15) | 0.50 | -1.85 | (-4.23, 0.54) | -0.04 | (-0.08, 0.01) | 0.50 |

Table 2. (continued)
|  |  |  |  |  |  |  |  |  |  |  |  |
| --- | --- | --- | --- | --- | --- | --- | --- | --- | --- | --- | --- |
| Cuculidae | 3 | -4.14 | (-4.97,<br>-3.36) | -0.42 | (-0.47,<br>-0.36) | 0.67 | -7.12 | (-8.74,<br>-5.52) | -0.39 | (-0.46,<br>-0.31) | 0.67 |
| Falconidae | 2 | 0.02 | (-0.01,<br>0.06) | 0.11 | (-0.04,<br>0.30) | 0.50 | -1.95 | (-2.29,<br>-1.63) | -0.41 | (-0.45,<br>-0.37) | 0.50 |
| Fringillidae | 5 | 4.85 | (3.29,<br>6.86) | 0.65 | (0.43,<br>0.93) | 0.60 | -67.84 | (-91.25,<br>-41.65) | -0.35 | (-0.44,<br>-0.22) | 0.80 |
| Hirundinidae | 6 | 14.81 | (9.08,<br>23.04) | 0.68 | (0.42,<br>1.07) | 0.33 | -36.59 | (-53.62,<br>-18.92) | -0.17 | (-0.24,<br>-0.09) | 0.83 |
| Icteridae | 9 | -119.28 | (-130.06,<br>-109.00) | -0.64 | (-0.66,<br>-0.62) | 0.89 | -260.66 | (-282.85,<br>-239.06) | -0.4 | (-0.42,<br>-0.38) | 0.89 |
| Icteriidae | 1 | -4.18 | (-5.40,<br>-3.07) | -0.28 | (-0.33,<br>-0.21) | 1.00 | -4.40 | (-6.12,<br>-2.73) | -0.24 | (-0.31,<br>-0.15) | 1.00 |
| Laniidae | 1 | -2.66 | (-3.15,<br>-2.19) | -0.80 | (-0.83,<br>-0.77) | 1.00 | -9.79 | (-11.17,<br>-8.48) | -0.69 | (-0.73,<br>-0.65) | 1.00 |
| Mimidae | 3 | -7.89 | (-9.37,<br>-6.50) | -0.23 | (-0.26,<br>-0.19) | 1.00 | -16.09 | (-20.34,<br>-11.82) | -0.19 | (-0.24,<br>-0.14) | 0.67 |
| Pandionidae | 1 | 0.06 | (0.03,<br>0.10) | 2.52 | (1.61,<br>3.73) | 0.00 | 0.31 | (0.23,<br>0.42) | 1.85 | (1.42,<br>2.34) | 0.00 |
| Paridae | 3 | 0.52 | (-0.49,<br>1.55) | 0.03 | (-0.03,<br>0.08) | 0.33 | 11.49 | (7.43,<br>15.85) | 0.23 | (0.15,<br>0.31) | 0.33 |
| Parulidae | 23 | -14.68 | (-17.49,<br>-11.78) | -0.19 | (-0.22,<br>-0.16) | 0.57 | -54.25 | (-67.36,<br>-40.79) | -0.14 | (-0.17,<br>-0.10) | 0.52 |
| Passerellidae | 11 | -48.38 | (-53.14,<br>-43.78) | -0.39 | (-0.42,<br>-0.36) | 0.73 | -439.63 | (-487.61,<br>-391.74) | -0.38 | (-0.41,<br>-0.34) | 0.91 |
| Picidae | 7 | 0.22 | (-0.61,<br>1.09) | 0.01 | (-0.03,<br>0.05) | 0.43 | 5.47 | (2.57,<br>8.71) | 0.09 | (0.04,<br>0.14) | 0.29 |
| Regulidae | 1 | 0.12 | (0.00,<br>0.45) | 0.74 | (0.07,<br>1.83) | 0.00 | -22.52 | (-45.76,<br>2.32) | -0.17 | (-0.33,<br>0.02) | 1.00 |
| Sittidae | 3 | 2.09 | (1.64,<br>2.60) | 0.78 | (0.60,<br>0.98) | 0.33 | 13.54 | (9.85,<br>18.19) | 0.84 | (0.61,<br>1.11) | 0.33 |
| Strigidae | 4 | 0.59 | (0.39,<br>0.82) | 0.27 | (0.18,<br>0.37) | 0.50 | 1.07 | (0.36,<br>1.81) | 0.13 | (0.04,<br>0.22) | 0.75 |
| Trochilidae | 1 | 4.28 | (2.79,<br>6.06) | 0.33 | (0.22,<br>0.45) | 0.00 | 12.50 | (9.54,<br>15.82) | 0.6 | (0.47,<br>0.75) | 0.00 |
| Troglodytidae | 4 | 5.44 | (3.86,<br>7.25) | 0.37 | (0.26,<br>0.49) | 0.25 | 26.93 | (19.80,<br>34.73) | 0.42 | (0.31,<br>0.54) | 0.00 |
| Turdidae | 5 | -3.67 | (-6.32,<br>-1.01) | -0.06 | (-0.11,<br>-0.02) | 0.40 | 13.96 | (-7.72,<br>36.05) | 0.03 | (-0.02,<br>0.08) | 0.40 |
| Tyrannidae | 9 | -5.44 | (-7.12,<br>-3.73) | -0.13 | (-0.17,<br>-0.09) | 0.44 | -71.82 | (-90.86,<br>-53.48) | -0.22 | (-0.27,<br>-0.17) | 0.89 |
| Tytonidae | 1 | 0.01 | (0.00,<br>0.03) | 2.73 | (1.72,<br>4.14) | 0.00 | 0.11 | (0.06,<br>0.19) | 1.98 | (1.32,<br>2.82) | 0.00 |
| Vireonidae | 6 | 14.67 | (12.24,<br>17.26) | 0.36 | (0.30,<br>0.43) | 0.17 | 73.72 | (62.23,<br>85.89) | 0.45 | (0.38,<br>0.53) | 0.00 |

**Fig. 4.**
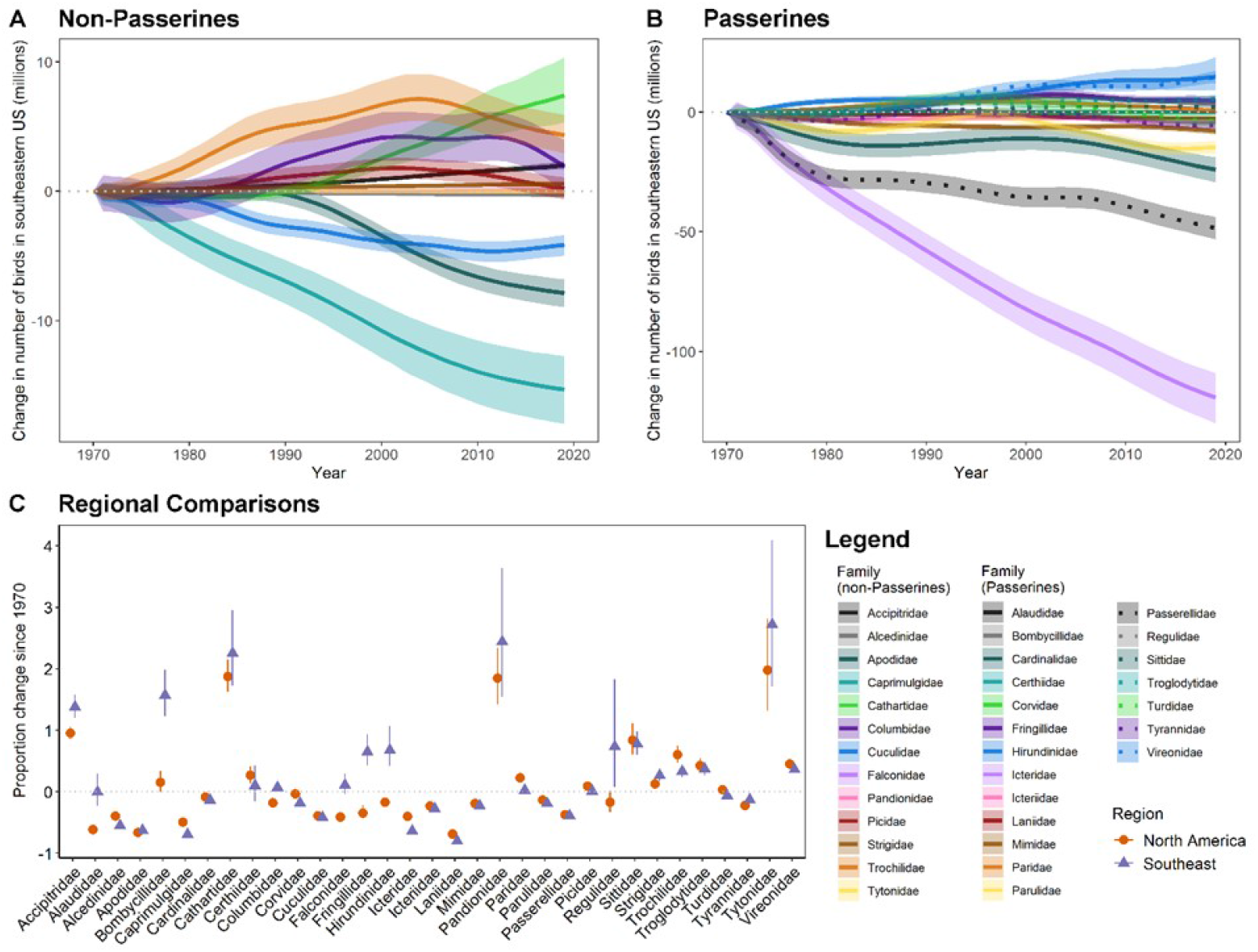
Net change in abundance of landbirds (n = 141 species) by family for (A) non-Passerines and (B) Passerines in the Southeastern United States, in millions. Comparisons of proportional change by family between the Southeast and North America are shown in (C). Annual median abundance change estimates (A, B) or overall proportion change (C) and 95% confidence intervals are shown.

Proportionally, Laniidae (Shrikes; n = 1 species [Loggerhead Shrike, *Lanius ludovicianus*]) exhibited the greatest relative population loss in the Southeast at -80% ([-0.83, -0.77]; Table 2). Two families of aerial insectivores also experienced large proportional declines in the region: Caprimulgidae declined by 69% (−0.72, -0.67), and Apodidae (Swifts, *n* = 1 species [Chimney Swift, *Chaetura pelagica*]) declined by 63% ([-0.65, -0.60]; Table 2). The largest proportional gains in the Southeast were of Tytonidae (Barn Owls; *n* = 1 species [Barn Owl, *Tyto alba*]) with a 273% increase (1.72, 4.14) and Pandionidae (Osprey; *n* = 1 species [Osprey, *Pandion haliaetus*]) with a 252% increase ([1.61, 3.73]; Table 2).

#### Southeast versus North America

Proportional losses among families in the Southeast were similar overall in comparison to North America (Fig. 4C). As in the Southeast, Laniidae had the greatest proportional losses in North America (−69% [-0.73, -0.65]), similarly so for Tytonidae with the largest proportional gain (North America 198%, [1.32, 2.82]). In total, 95% confidence intervals of Southeast and North American proportional change estimates overlapped for 18 of 33 families (54.55%; Fig. 4C).

### Habitat Association

#### Southeast

Among Level I habitat associations, the largest numeric losses in the Southeast were among birds associated with scrub-shrub / early seral habitat (*n* = 16 species), with a loss of -77.23 million birds ([-82.97, -71.67]; Table 3, Fig. 5A). Proportionally, the greatest losses in the Southeast were among grassland / open land birds (*n* = 21 species) at -54% ([-0.57, -0.51]) and emergent wetland birds (n = 6 species) at -51% (−0.54, -0.47). Conversely, birds using airspace for foraging (*n* = 7 species) increased in numeric abundance (+6.95 million birds, [1.06, 15.24]; Table 3, Fig. 5A), as well as yielding the greatest proportional increases in the Southeast (20% [0.03, 0.44]) while declining by -22% (−0.28, -0.14) across North America (Fig. 5B). The Southeast increase was strongly driven by an exponential 9,747% increase (68.56, 138.15) in Cliff Swallows (Appendix 1).

**Table 3.** Population change of native landbirds within Level-I habitat associations in the Southeastern US and the same species set across North America, 1970-2019. Number of species, abundance change (in millions), proportion abundance change, and proportion of declining species are shown for all native species excluding BGGN (n = 141).

| Level I Habitat Association | Number of Species | Southeastern US |  |  |  |  | North America |  |  |  |  |
| --- | --- | --- | --- | --- | --- | --- | --- | --- | --- | --- | --- |
|  |  | Numerical Abundance Change (Millions) |  | Proportion Abundance Change |  | Proportion Declining Species | Numerical Abundance Change (Millions) |  | Proportion Abundance Change |  | Proportion Declining Species |
|  |  | Change | 95% CI | Change | 95% CI |  | Change | 95% CI | Change | 95% CI |  |
| Aerial | 7 | 6.95 | (1.06, 15.24) | 0.20 | (0.03, 0.44) | 0.43 | -50.56 | (-67.66, -32.71) | -0.22 | (-0.28, -0.14) | 0.86 |
| Emergent wetland | 6 | -38.93 | (-45.26, -33.00) | -0.51 | (-0.54, -0.47) | 0.67 | -102.31 | (-119.88, -84.88) | -0.28 | (-0.32, -0.24) | 0.67 |
| Forest woodland | 79 | -3.41 | (-8.93, 2.12) | -0.01 | (-0.03, 0.01) | 0.42 | -124.77 | (-183.01, -66.61) | -0.06 | (-0.09, -0.03) | 0.44 |
| Generalist | 18 | -41.94 | (-50.81, -33.29) | -0.17 | (-0.20, -0.13) | 0.50 | -239.7 | (-267.66, -212.04) | -0.23 | (-0.26, -0.21) | 0.61 |
| Grassland / open land | 21 | -66.11 | (-73.20, -59.11) | -0.54 | (-0.57, -0.51) | 0.62 | -441.67 | (-479.45, -406.73) | -0.46 | (-0.48, -0.43) | 0.81 |
| Shrub-scrub / early successional | 16 | -77.23 | (-82.97, -71.67) | -0.34 | (-0.36, -0.32) | 0.75 | -124.81 | (-145.85, -105.16) | -0.22 | (-0.25, -0.19) | 0.69 |

**Fig. 5.**
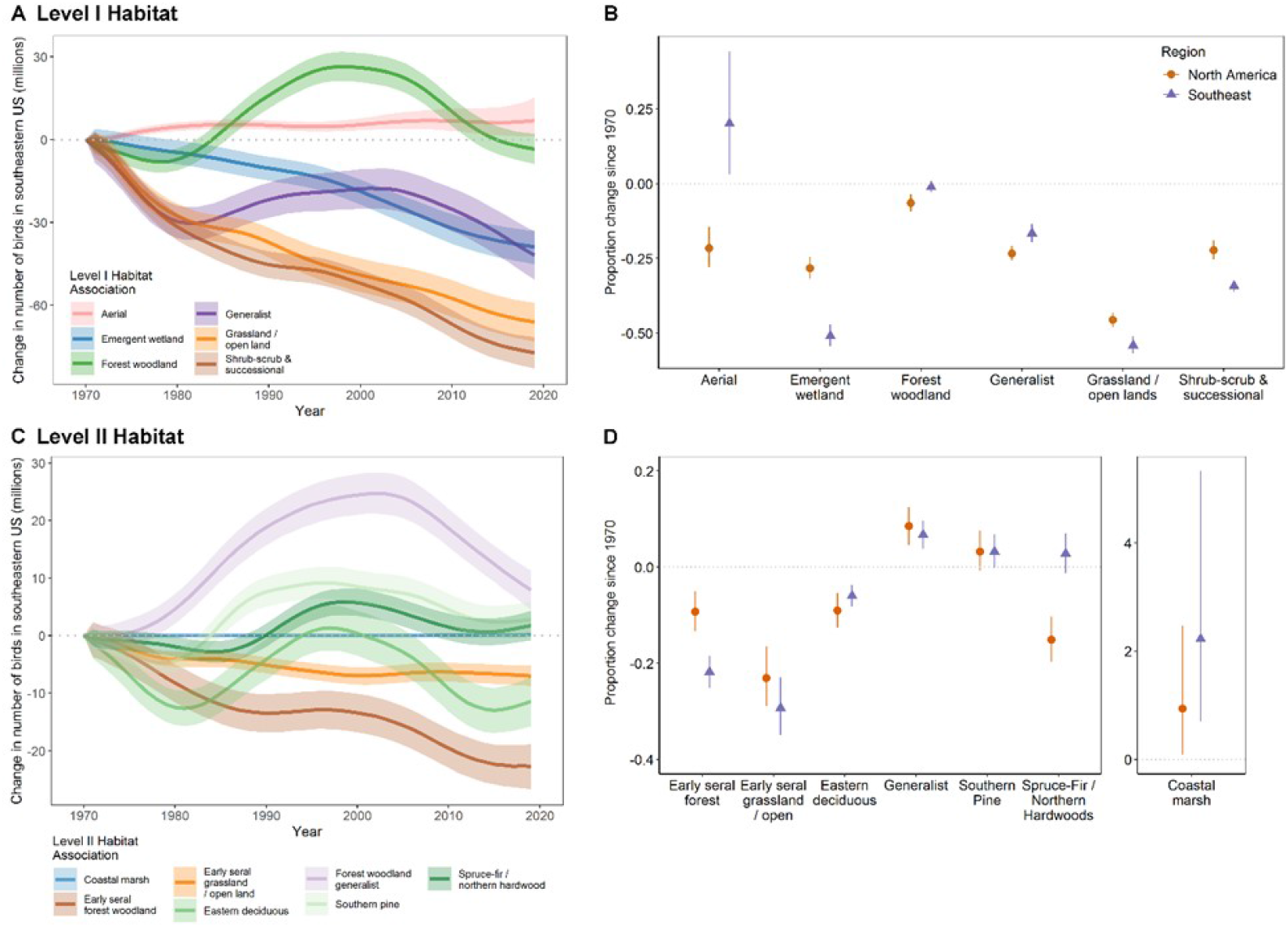
Net change in abundance of landbirds (n = 141 species) by (A) Level I and (B) Level II Habitat association in the Southeastern United States, in millions. Comparisons of proportional change by Level I (C) and Level II (D) Habitat association. Annual median abundance change estimates (A, C) or overall proportion change (B, D) and 95% confidence intervals are shown.

Three of the six Level I habitat associations experienced declines in the Southeast and North America, with greater proportional declines in the Southeast: emergent wetland, grassland / open lands, and shrub-scrub / early successional (Fig. 5B). The remaining three habitat associations (forest woodlands, aerial foragers, and generalists) declined less or increased more in the Southeast as compared to North America (Fig. 5B): all with non-overlapping 95% confidence intervals.

Among Level II habitat associations in the Southeast, the greatest numeric losses (−22.66 million birds, [-26.67, -18.79]) were among birds associated with early seral forest woodland (*n* = 7 species; Table 4, Fig. 5C). Proportionally, the greatest losses (−29% [-0.35, -0.23]) were of early seral grassland / open land birds (*n* = 2 species). Conversely, the greatest numeric gain was observed among forest woodland generalists (*n* = 20 species) which gained an estimated 7.96 million breeding birds (4.50, 11.33). Proportionally, birds in coastal marshes (*n* = 1 species [Seaside Sparrow, *Ammospiza maritima*]) had the greatest gains at an estimated 223% increase ([0.70, 5.33]; Table 4).

**Table 4.** Population change of native landbirds within Level-II habitat associations in the Southeastern US and the same species set across North America, 1970-2019. Number of species, abundance change (in millions), proportion abundance change, and proportion of declining species are shown for all native species excluding BGGN (n = 141).

| Level II Habitat Association | Number of Species | Southeastern US |  |  |  |  | North America |  |  |  |  |
| --- | --- | --- | --- | --- | --- | --- | --- | --- | --- | --- | --- |
|  |  | Numerical Abundance Change (Millions) |  | Proportion Abundance Change |  | Proportion Declining Species | Numerical Abundance Change (Millions) |  | Proportion Abundance Change |  | Proportion Declining Species |
|  |  | Change | 95% CI | Change | 95% CI |  | Change | 95% CI | Change | 95% CI |  |
| Coastal marsh | 1 | 0.14 | (0.00, 0.57) | 2.23 | (0.70, 5.33) | 0.00 | 0.10 | (0.00, 0.38) | 0.94 | (0.09, 2.47) | 0.00 |
| Early seral forest woodland | 7 | -22.66 | (-26.67, -18.79) | -0.22 | (-0.25, -0.18) | 0.86 | -16.40 | (-24.00, -8.74) | -0.09 | (-0.13, -0.05) | 0.71 |
| Early seral grassland / open land | 2 | -6.96 | (-8.72, -5.21) | -0.29 | (-0.35, -0.23) | 0.50 | -8.91 | (-11.63, -6.14) | -0.23 | (-0.29, -0.17) | 0.50 |
| Eastern deciduous | 51 | -11.41 | (-15.77, -6.99) | -0.06 | (-0.08, -0.04) | 0.43 | -98.52 | (-139.39, -58.45) | -0.09 | (-0.13, -0.05) | 0.49 |
| Forest woodland generalist | 20 | 7.96 | (4.50, 11.33) | 0.07 | (0.04, 0.10) | 0.35 | 46.55 | (25.33, 68.10) | 0.09 | (0.05, 0.12) | 0.25 |
| Southern pine | 17 | 2.81 | (-0.19, 5.94) | 0.03 | (0.00, 0.07) | 0.41 | 4.42 | (-1.08, 10.32) | 0.03 | (-0.01, 0.08) | 0.47 |
| Spruce-fir / northern hardwood | 31 | 1.78 | (-0.80, 4.35) | 0.03 | (-0.01, 0.07) | 0.39 | -156.73 | (-208.98, -105.44) | -0.15 | (-0.20, -0.10) | 0.45 |

#### Southeast versus North America

Early seral grassland / open land birds experienced the greatest proportional losses across North America (−23% [-0.29, -0.17]) and in the Southeast (Table 4; Fig. 5D). Conversely, coastal marsh birds experienced the greatest proportional increases across North America (94% [0.09, 2.47]) and in the Southeast. Birds that associate, at least in part, with spruce-fir / northern hardwoods habitat increased slightly in the Southeast (3% [-0.01, 0.07]) despite declining across North America (−15% [-0.20, -0.10]). Conversely, early seral forest birds declined more in the Southeast (−22% [-0.25, -0.18]) than across North America (−9% [-0.13, -0.05]). All other Level II habitat associations (*n* = 5) experienced similar gains or losses in both regions (Fig. 5D).

### Migratory Status

#### Southeast

In the Southeast, the greatest numeric losses were among partial migrants (*n* = 50), which saw an estimated loss of -151.12 million birds ([-163.77, -138.75]; Table 5, Fig. 6A). Partial migrants also experienced the greatest proportional losses, at -30% (−0.32, -0.28). The greatest numeric gains (8.18 million birds [3.42, 13.02]) and proportional gains (6% [0.02, 0.10]) were observed among resident species (*n* = 25 species).

**Table 5.** Population change of native landbirds by migratory status in the Southeastern US and the same species set across North America, 1970-2019. Number of species, abundance change (in millions), proportion abundance change, and proportion of declining species are shown for all native species excluding BGGN (n = 141).

| Migratory Status | Number of Species | Southeastern US |  |  |  |  | North America |  |  |  |  |
| --- | --- | --- | --- | --- | --- | --- | --- | --- | --- | --- | --- |
|  |  | Numerical Abundance Change (Millions) |  | Proportion Abundance Change |  | Proportion Declining Species | Numerical Abundance Change (Millions) |  | Proportion Abundance Change |  | Proportion Declining Species |
|  |  | Change | 95% CI | Change | 95% CI |  | Change | 95% CI | Change | 95% CI |  |
| Migrant | 66 | -56.31 | (-64.91, -46.32) | -0.18 | (-0.20, -0.15) | 0.53 | -285.35 | (-327.09, -244.6) | -0.17 | (-0.20, -0.15) | 0.64 |
| Partial migrant | 50 | -151.12 | (-163.77, -138.75) | -0.30 | (-0.32, -0.28) | 0.50 | -801.58 | (-870.33, -734.01) | -0.27 | (-0.29, -0.25) | 0.58 |
| Resident | 25 | 8.18 | (3.42, 13.02) | 0.06 | (0.02, 0.10) | 0.40 | 45.29 | (35.35, 55.60) | 0.17 | (0.13, 0.21) | 0.36 |

**Fig. 6.**
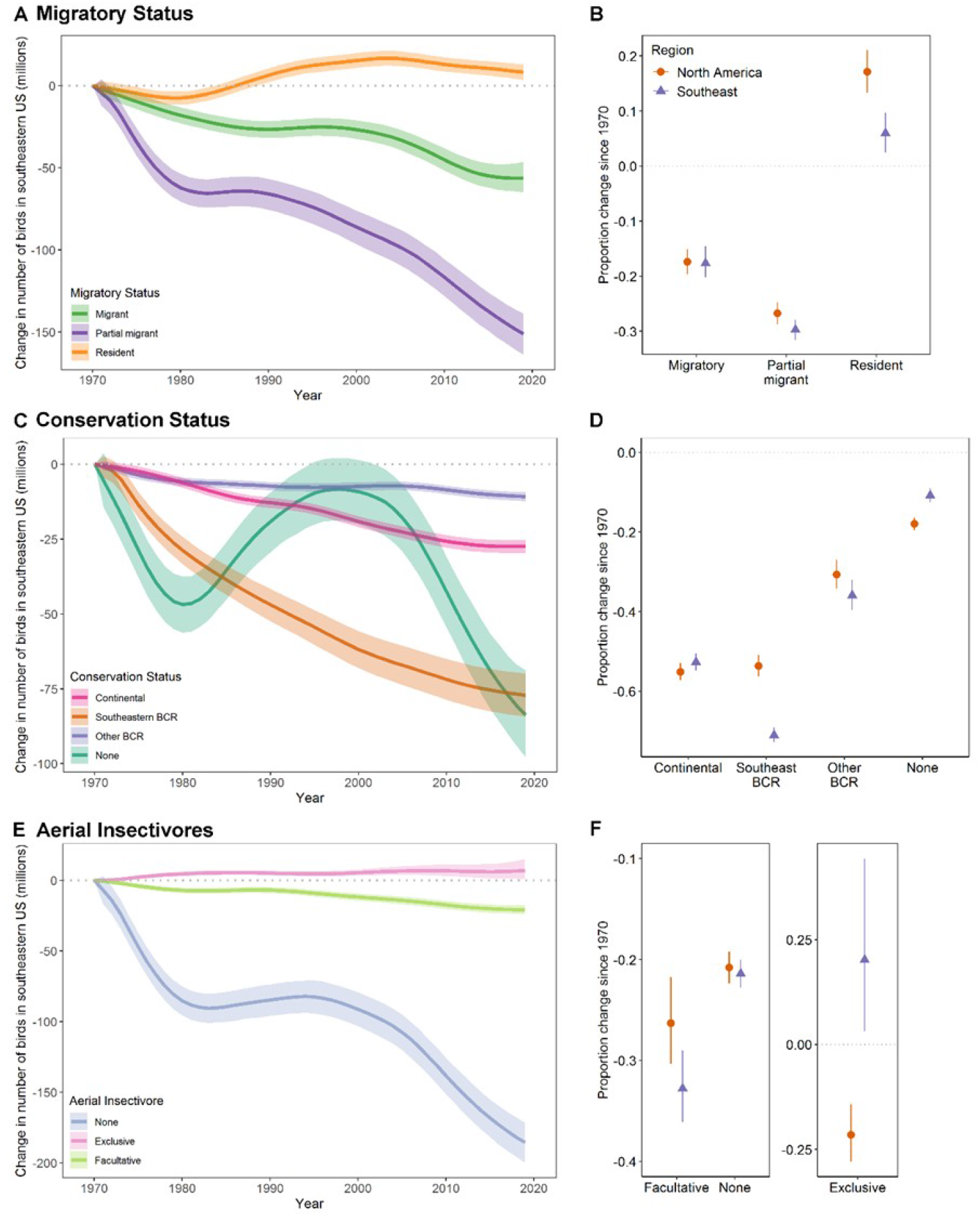
Net change in abundance of landbirds (n = 141 species) by (A) Migratory Status, (C) Conservation Status, and (E) Aerial Insectivores group in the Southeastern United States, in millions. Comparisons of proportional change are shown in (B, D, F). Annual median abundance change estimates (A, C, E) or overall proportion change (B, D, F) and 95% confidence intervals are shown.

#### Southeast versus North America

In comparison, partial migrants also experienced the greatest proportional losses across North America (−27%, [-0.29, -0.25]; Table 5; Fig. 6B) and resident birds experienced the greatest proportional increases across North America (17% [0.13, 0.21]). Numerically, resident species increased more across North America than in the Southeast (Fig. 6B).

### Conservation Status

#### Southeast

Species identified as Birds of Conservation Concern (BCC; *n* = 33 species) lost a total of -115.42 million birds ([-126.39, -104.44]; Table 6, Fig. 6C) in the Southeast. Those species listed as BCCs in the Southeast at only the BCR level (*n* = 13 species) exhibited the greatest numeric losses (−77.16 million birds [-84.31, -69.85]), followed by species (*n* = 20) listed as BCCs at the continental scale (−27.44 million birds [-29.66, -25.31]). Proportionally, species listed as BCCs in the Southeast only at the BCR level lost -71% (−0.73, -0.69) of their 1970 population, whereas continentally listed BCC species lost -53% (−0.55, -0.51). In comparison, species not listed as BCCs (*n* = 108 species) lost -11% (−0.13, -0.09) of their 1970 population

**Table 6.**
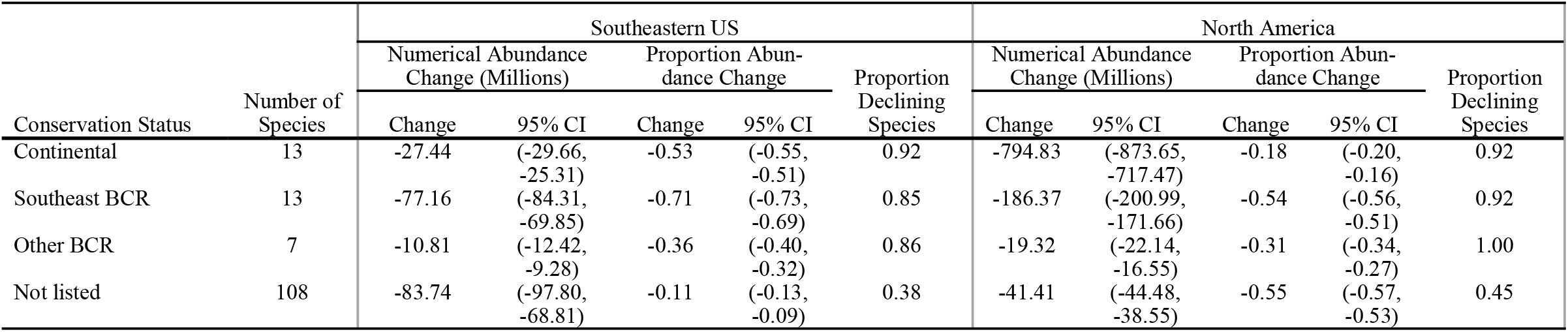
Population change of native landbirds by conservation status in the Southeastern US and the same species set across North America, 1970-2019. Number of spe cíes, abundance change (in millions), proportion abundance change, and proportion of declining species are shown for all native species excluding BGGN (n = 141).

#### Southeast versus North America

Across North America, continentally listed BCC species exhibited the greatest proportional losses (−55% [-0.57, -0.53]; Table 6; Fig. 6D), whereas in the Southeast the greatest losses were among species listed as BCCs at the BCR level only. Species listed as BCC in Southeastern BCRs declined proportionally more in the Southeast than across North America (−56% [-0.56, -0.51]). Conversely, species not listed as BCC declined more across North America (−18% [-0.20, -0.16]) than the Southeast. Continental BCC species and species listed for BCRs outside the Southeast (n = 7 species) experienced similar gains or losses in both regions (Fig. 6D).

### Aerial Insectivores

#### Southeast

Aerial insectivores in the Southeast (*n* = 19 species) lost -13.83 million birds (−22.87, -2.29), but these losses primarily occurred among 12 species of facultative aerial insectivores (i.e., species that do not strictly forage aerially; Table 7, Fig. 6E). These facultative species lost -20.78 million birds (−23.93, -17.53) or -33% (−0.36, -0.29) of their 1970 population. Conversely, obligate aerial insectivores (n = 7) in the Southeast increased by an estimated 6.95 million birds (1.06, 15.24) or 20% (0.03, 0.44) since 1970. Four of the seven obligate aerial insectivores increased in abundance, including an 805% increase in Tree Swallows (*Tachycineta bicolor*) and a 9,747% increase in Cliff Swallows. Three species of obligate aerial insectivores declined consistent with continental patterns in this guild, including a - 76% decline in Bank Swallows (Riparia riparia) and a -63% decline in Chimney Swifts (Fig. 6E, Appendix 1).

**Table 7.** Population change of aerial insectivores in the Southeastern US and the same species set across North America, 1970-2019. Number of species, abundance change (in millions), proportion abundance change, and proportion of declining species are shown for all native species excluding BGGN (n = 141).

| Aerial Insectivores | Number of Species | Southeastern US |  |  |  |  | North America |  |  |  |  |
| --- | --- | --- | --- | --- | --- | --- | --- | --- | --- | --- | --- |
|  |  | Numerical Abundance Change (Millions) |  | Proportion Abundance Change |  | Proportion Declining Species | Numerical Abundance Change (Millions) |  | Proportion Abundance Change |  | Proportion Declining Species |
|  |  | Change | 95% CI | Change | 95% CI |  | Change | 95% CI | Change | 95% CI |  |
| Exclusive aerial insectivore | 7 | 6.95 | (1.06, 15.24) | 0.20 | (0.03, 0.44) | 0.43 | -50.56 | (-67.66, -32.71) | -0.22 | (-0.28, -0.14) | 0.86 |
| Facultative aerial insectivore | 12 | -20.78 | (-23.93, -17.53) | -0.33 | (-0.36, -0.29) | 0.58 | -99.12 | (-118.44, -80.14) | -0.26 | (-0.30, -0.22) | 0.92 |
| Neither | 122 | -185.51 | (-199.91, -171.55) | -0.21 | (-0.23, -0.20) | 0.49 | -891.68 | (-967.80, -817.84) | -0.21 | (-0.22, -0.19) | 0.52 |

#### Southeast versus North America

In comparison to estimates across North America, facultative aerial insectivores also experienced the greatest proportional losses (−26% [-0.30, -0.22]; Table 7; Fig. 6F). Whereas obligate aerial insectivores declined more across North America (−22% [-0.28, -0.14]) than in the Southeast, non-aerial insectivores (*n* = 122 species) experienced similar gains or losses in both regions (Fig. 6F).

### Bird Conservation Regions

Across the Southeast, bird populations declined in all BCRs with net abundance change ranging from -59.72 million birds (−65.71, -53.07) in the Appalachian Mountains (BCR 28, *n* = 125 species), to -3.18 million birds (−10.93, 40.28) in the Mississippi Alluvial Valley (BCR 26, *n* = 99 species; Table 8, Fig. 7A-H). However, because the eight BCRs in the Southeast region varied greatly in size (Fig. 1) and number of breeding species included in the analysis (Table 8), proportional change estimates provide the most reliable comparisons among BCRs. Proportionally, the greatest losses by far occurred in Peninsular Florida (BCR 31, *n* = 77 species) at -60% (−0.68, -0.48) followed by the Gulf Coastal Prairie (BCR 37, *n* = 9 species) at -29% ([-0.38, -0.16]; Fig. 7I). Peninsular Florida also had the largest proportion of declining species (67%), while proportions of declining species in other BCRs ranged between 41-57% (Table 3). No proportional gains were observed in any BCR, but the smallest declines occurred in the Mississippi Alluvial Valley with a -6% loss (−0.19, 0.70).

**Table 8.** Population change of native landbirds by Bird Conservation Region in the Southeastern US, 1070-2019. Number of species, abundance change (in millions), proportion abundance change, and proportion of declining species are shown for all native species excluding BGGN (n =141). See Appendix 3 for Group-level summaries by BCR.

| Bird Conservation Region (BCR) |  | Number of Species | Numerical Abundance Change (Millions) |  | Proportion Abundance Change |  | Proportion Declining Species |
| --- | --- | --- | --- | --- | --- | --- | --- |
| BCR Code | BCR Name |  | Change | 95% CI | Change | 95% CI |  |
| 24 | Central Hardwoods | 110 | -33.62 | (-40.77, -25.58) | -0.19 | (-0.23, -0.15) | 0.45 |
| 25 | West Gulf Coastal Plain | 104 | -21.82 | (-28.2, -13.10) | -0.22 | (-0.28, -0.13) | 0.57 |
| 26 | Mississippi Alluvial Valley | 99 | -3.18 | (-10.93, 40.28) | -0.06 | (-0.19, 0.70) | 0.43 |
| 27 | Southeastern Coastal Plain | 108 | -36.04 | (-45.77, -5.00) | -0.16 | (-0.19, -0.02) | 0.49 |
| 28 | Appalachian Mountains | 125 | -59.72 | (-65.71, -53.07) | -0.23 | (-0.25, -0.21) | 0.53 |
| 29 | Piedmont | 110 | -13.32 | (-17.00, -9.46) | -0.14 | (-0.18, -0.10) | 0.46 |
| 31 | Peninsular Florida | 75 | -33.13 | (-47.87, -20.34) | -0.60 | (-0.68, -0.48) | 0.67 |
| 37 | Gulf Coastal Prairie | 86 | -8.25 | (-12.67, -3.92) | -0.29 | (-0.38, -0.16) | 0.41 |

**Fig. 7.**
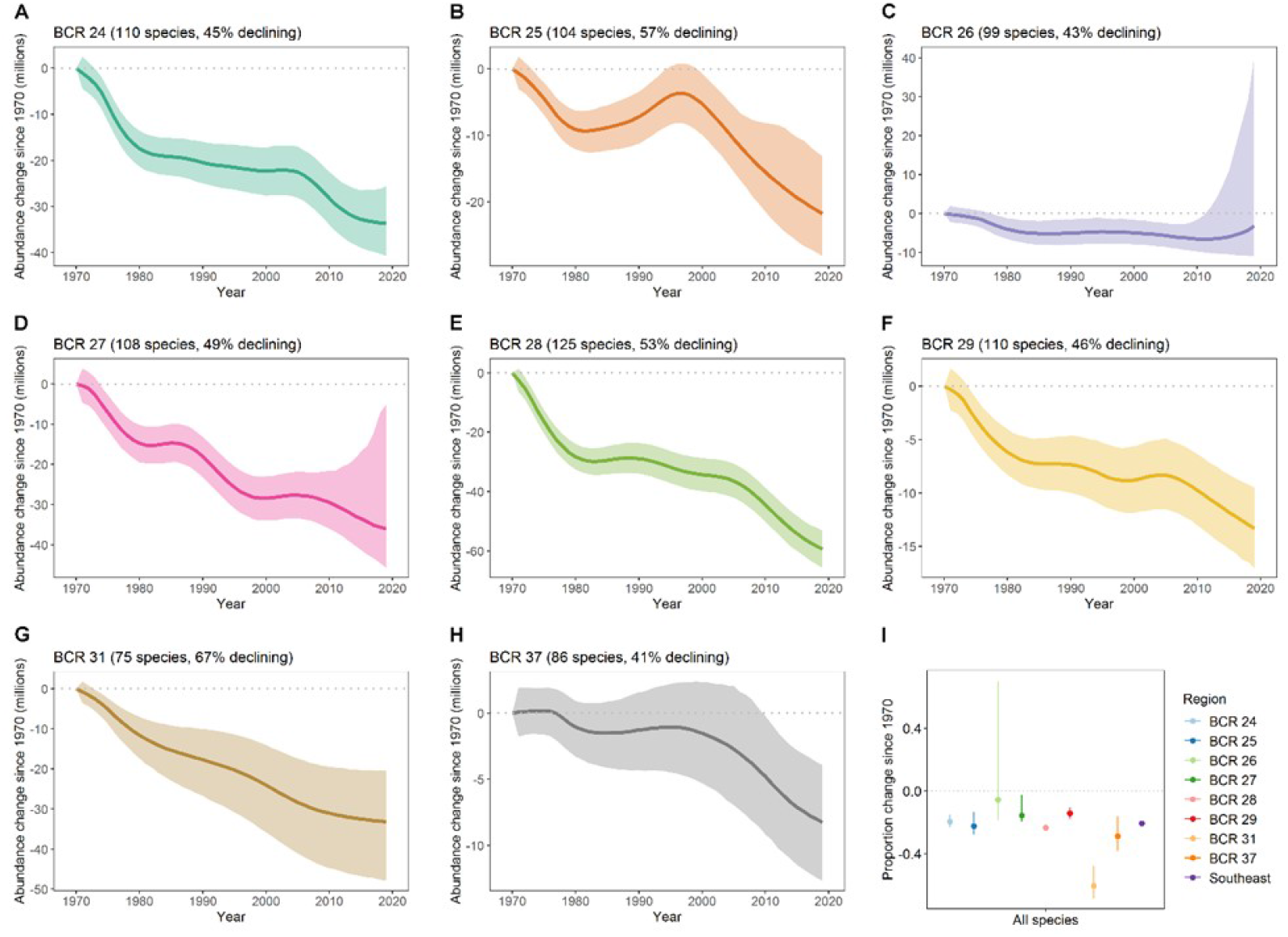
Net change in abundance of landbirds (n = 141 species) by Bird Conservation Region (BCR). Annual median abundance change estimates and 95% confidence intervals are shown.

#### Population Change by Habitat Association within BCRs

Population changes among Level I habitat association groups was generally similar among BCRs Appendix 3; Fig. 8A), though species using air space (i.e., aerial insectivores) proportionally increased more in the Mississippi Alluvial Valley. Generalists and shrub-scrub / early successional birds increased more in the Gulf Coast Prairies. Grassland / open land and shrub-scrub / early seral birds decreased more in Peninsular Florida. Level II habitat association groups also showed broadly similar population change (Appendix 3; Fig. 8B), though early seral forest woodland birds increased only in Peninsular Florida and the Gulf Coast Prairies. Southern pine birds decreased more and spruce-fir & northern hardwood birds increased more in Peninsular Florida and coastal marsh and early seral grassland / open land birds increased more in the Gulf Coast Prairies.

**Fig. 8.**
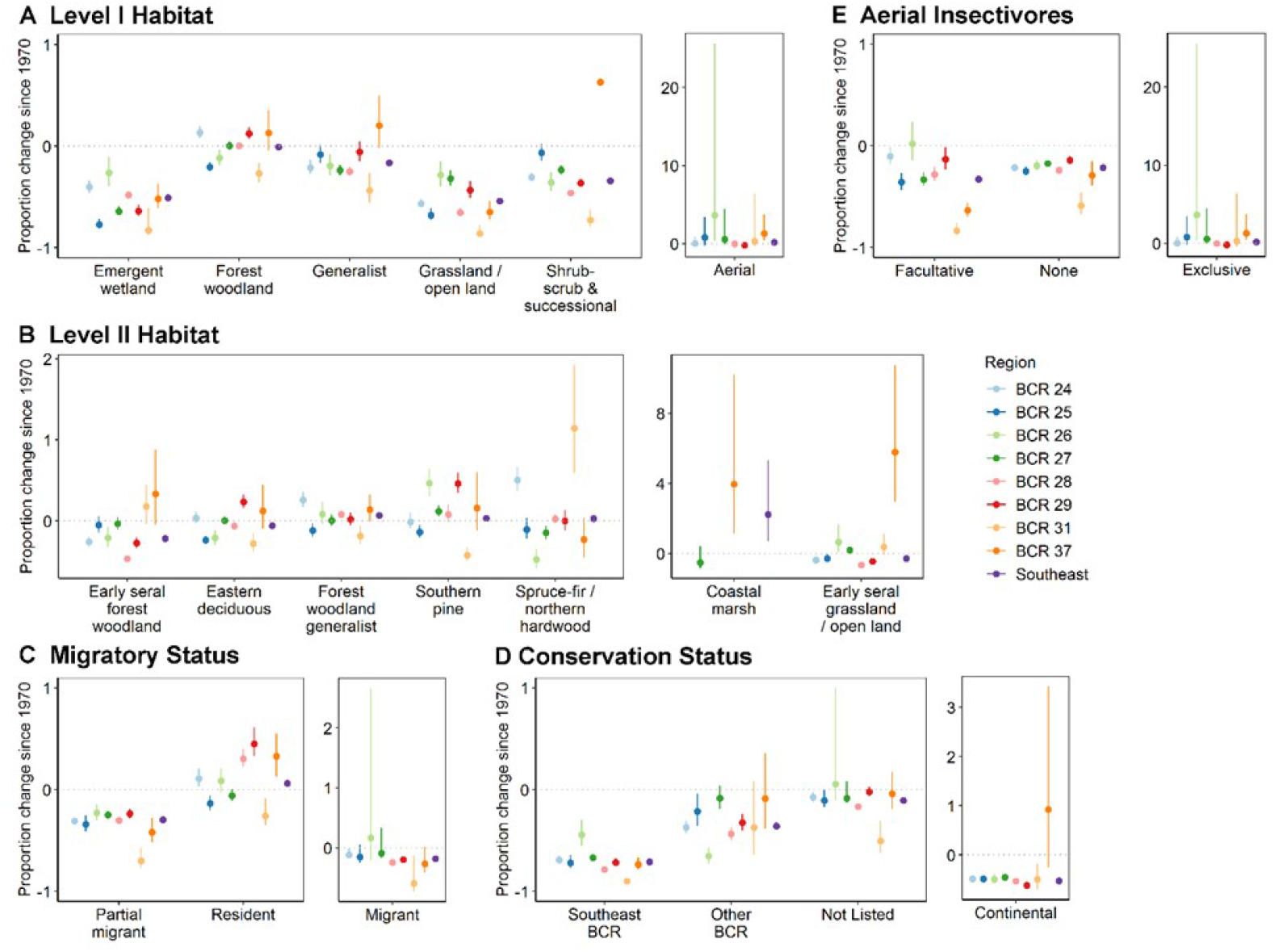
Comparisons of proportional change in abundance of landbirds (n = 141 species) between the Southeast Region and each of the eight BCRs included in the region. Overall proportion change and 95% confidence intervals are shown.

#### Population Change by Migratory Status within BCRs

The largest proportional losses of all three migratory status categories occurred in Peninsular Florida (Appendix 3). Proportionally, resident species increased more in the Piedmont (BCR 29), while migrants increased more in the Mississippi Alluvial Valley. Proportional loss for partial migrants was similar across the Southeast (Fig. 8C), except for Peninsular Florida showing significantly greater loss (i.e., non-overlapping confidence intervals).

#### Population Change by Conservation Status within BCRs

Continentally listed BCC species appeared to increase in the Gulf Coast Prairies though the 95% confidence interval included zero, unlike the other BCRs where this group of species declined (Appendix 3; Fig. 8D). Species listed as BCC for one or more Southeastern BCRs declined less in the Mississippi Alluvial Valley, while species listed for one or more non-Southeastern BCRs declined more in Peninsular Florida.

#### Population Change by Aerial Insectivores within BCRs

Declines in facultative aerial foragers were greater in Peninsular Florida and the Gulf Coast Prairies. Whereas the smallest declines were observed in the Mississippi Alluvial Valley. Obligate aerial insectivores increased most strongly in the Mississippi Alluvial Valley, while populations of this guild remained generally stable elsewhere across the Southeast (Appendix 3; Fig. 8E).

## DISCUSSION

### Southeast versus North America

Rosenberg et al. (2019) documented the loss of nearly three billion birds across North America. Here, we assessed changes in landbird populations across the Southeast using the same analytical methods as described by Rosenberg et al. (2019) with a few deviations to account for data limitations (e.g., lack of population estimates for some species; see Methods). We also constrained our analyses to BCRs that overlap the USFWS’s Southeast Region. Our initial analyses (*n* = 146 bird species) showed a loss of 260 million birds since the 1970s. This represents a -22% loss, somewhat smaller than the -29% loss that occurred across North America (Rosenberg et al. 2019). Upon review of this initial assessment, it was determined that the Blue-gray Gnatcatcher comprised 16% of data, thereby potentially inserting undue influence on group-level analyses (e.g., habitats, migratory status, etc.). As such, we removed this species from all group-level analyses. Further, because we desired to evaluate bird species in a manner to inform conservation decision making and potential investment of resources (e.g., personnel, time, and funding), we also removed all non-native bird species (i.e., Eurasian Collared-Dove, European Starling, House Sparrow, and Rock Pigeon) resulting in a sample size of 141 bird species for group-level assessments. We found that roughly half of the 141 native landbird species declined since the 1970s, with total losses approaching 200 million birds.

Though landbirds declined overall in a similar proportion across the Southeast and North America since the 1970s (Table 1). Population losses were neither universal nor uniform within the Southeast, nor were such losses similar when compared to patterns in North America for relevant groupings of species (Tables 2-8). All 5 species groups (taxonomic family, habitat, migration status, BCC status and aerial foraging behavior) analyzed herein had at least one category that showed significant differences in proportional change over the 1970-2019 period between the Southeast and North America. These nuances contradict our null hypothesis (no difference between Southeast and North America rates of population change) and further support our assumption that these species groupings are relevant to conservation planning and implementation. Our findings also highlight important variation in population changes over the course of 1970-2019. However, our findings also highlight the need to carefully identify relevant groupings and to produce actionable results.

### Taxonomic Family

In the Southeast, 12 families fared worse or improved less when compared to North American estimates of population change. Alcedinidae (Belted Kingfisher, *Megaceryle alcyon*), Nightjars (Caprimulgidae), Cardinals and allies (Cardinalidae), Crows and Jays (Corvidae), Blackbirds (Icteridae), Shrikes (Laniidae), and Thrushes (Turdidae) decreased more in the Southeast than across North America; while Treecreepers (Certhiidae), Chickadees and Titmice (Paridae), Woodpeckers (Picidae), Hummingbirds (Trochilidae), and Vireos (Vireonidae) increased less in the Southeast as compared to North America. The superlative losses among these groups both regionally and continentally suggest pervasive problems affecting these groups at broad geographic scales.

Conversely, nine families fared better in the Southeast compared to North American estimates of population change. Larks (Alaudidae), Pigeons and Doves (Columbidae), Falcons (Falconidae), Finches (Fringillidae), Swallows (Hirundinidae), and Kinglets (Regulidae) all remained stable or increased in the Southeast despite declines at the North American scale. Flycatchers (Tyrannidae) declined in both regions but declined proportionally less in the Southeast. Conversely, Hawks (Accipitridae) and Cedar Waxwing (Bombycillidae; *Bombycilla cedrorum*) increased both continentally and in the Southeast but increased proportionally more in the Southeast. Other families, including Barn Owl (Tytonidae), Osprey (Pandionidae), and Vultures (Cathartidae) increased dramatically in both regions.

In drawing comparisons with patterns of gain or loss as reported by Rosenberg et al. (2019) for North America, it is important to note that species composition of most taxonomic categories (i.e., families) in the Southeast differed as a function of a more limited pool of regionally breeding species. Thus, while comparison of Regulidae between the two spatial-scales represents a comparison intrinsic to the same two species (Ruby -crowned and Golden-crowned Kinglets, Regulus calendula and R. satrapa, respectively), comparison of Picidae (woodpeckers), Mimidae (mockingbirds and thrashers), Sittidae (Nuthatches), Strigidae (true owls) and many others involve vastly different number of species within a given taxonomic group at the two spatial scales. Inferences or explanations regarding similar or different patterns in a taxonomic group at regional and continental scales must therefore take into consideration whether uniqueness in species composition may be an important factor in the observed pattern.

Still, gross comparisons revealing consistent patterns of gain or loss at both the regional and continental level may further highlight or reinforce taxonomic groups that appear to be particularly vulnerable (or adaptable). With this in mind, we constrained North American comparisons for group-level assessments to the same suite of species (i.e., species included in North America estimates are limited to the same suite of species used for Southeast estimates).

The similar number of families faring better or worse in the Southeast in comparison to North America highlights the idiosyncratic nature of how ecological processes impact bird populations when viewed at scale across multiple species (see Michel et al. 2021). To better understand these patterns of change requires in-depth analyses by a variety of species groupings at finer resolutions (e.g., local geographies, habitat, migratory status, foraging behavior, etc.).

### Habitat Associations

Proportional population changes among Level I habitat associations suggested that forest woodland birds, aerial foragers, and habitat generalists fared better in the Southeast than across North America. Aerial foragers increased in the Southeast while declining across North America, while the other two groups declined less regionally than continentally. However, birds associated with emergent wetland habitats declined nearly two times as much in the Southeast, and grassland / open land and shrub-scrub / early successional species also declined more in the Southeast when compared to North America estimates of population change.

#### Forest Woodlands

The relatively stable population trends of forest woodland birds across the Southeast is encouraging given the recent declines in the mid-2000s (see Fig. 5A and 5C). That is, starting in the mid -1980s forest woodland birds and habitat generalists across the Southeast underwent a population increase that lasted till the mid-2000s, whereas grassland / open land and shrub-scrub and early successional bird populations continued a downward population trajectory. Further evaluation of Level II habitats suggested this same relatively stable pattern held for eastern deciduous, southern pine, spruce-fir / northern hardwood birds and forest woodland generalists. Plausible explanations for this population increase include an increase in food resources (e.g., insect outbreak) and/or increased number of BBS routes (i.e., the mid-1990 time period coincides with the early beginnings of Partners in Fight and increased attention on landbirds). However, one would expect to see similar population increases across all habitats if either mechanism was the driving force. Further, the same pattern held after we removed Blue-gray Gnatcatcher, suggesting that a single, super abundant species was not driving the observed patterns. As such, we are left to accept the pattern as an unexplained phenomenon until further investigations can be conducted.

#### Aerial

Aerial insectivorous birds also exhibited an increasing population trajectory in the Southeast relative to North America. Proportional loss of facultative aerial insectivores (i.e., species that forage aerially, but not exclusively) in the Southeast was slightly greater (−33%) than for North America (−26%). However, obligate aerial insectivores – including swifts, swallows, and martins that forage exclusively in the air column, and which have been documented as undergoing extensive declines in other regions – increased by 20% in the Southeast. Dramatic increases in Cliff Swallows and Tree Swallows in the Southeast appear to be driving this regional pattern. While Cliff Swallow also increased across North America (+29 million birds), this was partially offset by continental losses in Tree Swallow estimated at -8.1 million birds. Growth in Cliff Swallow populations continent-wide appears to be a function of regional population growth in the Midwest and Southeast (Brown et al. 2020, Sauer et al. 2020), suggesting that increases in this species may have had a more pronounced influence on the overall pattern for obligate aerial insectivores in the Southeast. A recent study found that swallows were heavily impacted by climate indices (e.g., El Niño, North Atlantic Oscillation) and winds and storms during migration, particularly in western North America (Michel et al. 2021). This suggests that these species’ populations may be less susceptible to certain large-scale climatological impacts in the Southeast relative to other regions of North America.

#### Emergent Wetlands

Emergent wetlands and the birds that use them, are under threat in the Southeast. Across the U.S., over half of wetland area has been lost since European settlement, primarily to draining for agriculture and forestry (Mitsch and Gosselink 2007). The Southeast has been particularly hard hit, accounting for >85% of national wetland losses between the 1950s-1980s (Kurki-Fox et al. 2022). Not surprisingly, our results suggest bird populations associated with emergent wetlands have also declined. Of the emergent wetland species analyzed here, Red-winged Blackbird (*Ageliaius phoeniceus*) and Common Yellowthroat (*G. trichas*) contributed significantly (−27 million birds and -14 million birds, respectively) to the declines in emergent wetland bird communities across the Southeast. Red-winged Blackbird declines have been attributed largely to loss and degradation of emergent marsh habitat due to agricultural practices (Yasukawa and Searcy 2020), while Common Yellowthroat declines are less well-studied, but likely also related to habitat loss and fragmentation (Guzy and Ritchison 2020).

#### Grasslands / Open Lands

Grassland birds face multiple threats including habitat loss on both their breeding and wintering grounds across their range (Pool et al. 2014, Stanton et al. 2018, Wilsey et al. 2019). Over 90% of grassland habitats have been lost across the Southeast (Noss et al. 2021). Not surprisingly, grassland / open land birds showed steep, linear population declines across the Southeast; even out-pacing population losses at the scale of North America. Losses within this grassland / open land habitat were led by Eastern Meadowlark, Loggerhead Shrike (*L. ludovicianus*) and Grasshopper Sparrow (*Ammodramus savannarum*); all with >80% loses of their 1970s population. These alarming declines underpin the urgency for landscape-scale coordination and collaboration to protect and conserve remaining grassland habitats and restore degraded grassland habitats (to include lands converted to rowcrop agriculture) in an attempt to stabilize (or recover) negative population trends of associated bird populations. Collaborative, multi-partner efforts such as the Central Grasslands Roadmap (https://www.grasslandsroadmap.org/) are urgently needed in the Southeast and continentally to conserve these highly vulnerable species. Initiatives such as Audubon’s Conservation Ranching program that partner with private landowners to manage habitat for grassland birds (Michel et al. 2020) are particularly relevant, as the vast majority (84%) of remaining grasslands are in private ownership (North American Bird Conservation Initiative, U.S. Committee 2011).

#### Scrub-shrub / Early Seral

Similarly, the extent of shrub-scrub and early seral habitats continue to decline across North America with population trends of birds associated with these habitats following a similar decline across the Southeast (King and Schlossberg 2014). Our analyses showed that species of birds associated with these habitats lost a greater proportion of their population in the Southeast compared to North America. More specifically, analyses of Level II habitats showed that the Southeast lost a significantly greater proportion of early-seral forest birds when compared to North America estimates. This is especially alarming for species such as the Golden-winged Warbler, that lost 97% of its breeding population in the Southeast since the 1970s. Historical disturbance regimes such as wind-throw, fire, and flooding have largely been suppressed or altered by anthropogenic activities. King and Schlossberg (2014) suggested that conservation of these bird communities would require dedicated management actions and effort that focused on the ephemeral, early seral conditions.

### Migratory Status

Whereas migratory species experienced losses across North America (−17% for migrants and -27% for partial migrants), resident species increased by 17%. Similarly, we found the greatest proportional losses in the Southeast for species categorized as partial migrants (−30%), with lesser proportions lost for species categorized as long-distance migrants (−18%), with residents increasing by 6%. These results are contrary to findings in Europe which indicated that birds with partial migration strategies (i.e., overlapping breeding and non-breeding ranges) were less likely to experience population declines (Gilroy et al. 2016). One plausible explanation could be related to climate change. Recent research by Michel et al. (2021) found that among 37 species of aerial insectivores, climate impacts were strongest on short distance migrants (i.e., those that winter in the southern U.S., Mexico, and the Caribbean). Species that winter in more temperate latitudes may be experiencing more severe or pervasive negative effects from climate change than birds wintering in more tropical latitudes, thus contributing to observed differences in declines among these migratory groups. However, population increases of resident species runs counter to the “climate change” hypothesis”, in that, one would expect to observe population declines (not increases) for resident species if climate change was the single causal mechanism influencing their populations. Thus, additional full annual life cycle analyses are warranted to isolate factors affecting populations across seasons (i.e., breeding season, wintering season, and migration).

### Conservation Status

The USFWS recently updated the Birds of Conservation Concern (BCC) list which highlights species of nongame birds that without further conservation actions are likely to become candidates for listing under the U.S. Endangered Species Act (U.S. Fish and Wildlife Service 2021). Because BCCs are already recognized in-part due to declining populations, it’s unfortunate, but not surprising that populations of BCC birds exhibited greater proportional losses than non-BCC species. The greatest proportional losses were among the 13 species designated as BCCs at the level of a Southeastern BCR (−71%), followed by BCCs designated at the continental level (−53%). Among individual BCC species, the greatest numerical losses were estimated for Eastern Meadowlark (−40 million birds), a BCC in the West Gulf Coastal Plain / Ouachitas. Other substantial numerical losses among BCCs were estimated for Field Sparrow (*Spizella pusilla*, -10 million birds; a BCC in the Central Hardwoods), and Wood Thrush (−9 million birds; a continental level BCC). These strongly declining species each have different habitat associations and migratory statuses, highlighting the complexity of bird population declines, drivers, and consequently, the management actions needed to slow or reverse these declines.

Though we did not explicitly analyze them as a group here, our species list included five species that are included on the Road to Recovery’s Tipping Point list (North American Bird Conservation Initiative 2022). Tipping Point species include 70 species that have lost atleast 50% of their populations since 1970 and are predicted to lose an additional 50% of their current populations over the next 50 years without conservation action. Southeastern Tipping Point species include Chimney Swift, Golden-winged Warbler, Prairie Warbler (*Setophaga discolor*), Henslow’s Sparrow (*Centronyx henslowii*), and Bachman’s Sparrow (*Peucaea aestivalis*). All five of these species are also listed continentally as BCC (U.S. Fish and Wildlife Service 2021), indicating the severity of their respective population declines.

### Bird Conservation Regions

In addition to variation in bird population losses between the Southeast and other regions of North America (Rosenberg et al, 2019, North American Bird Conservation Initiative 2022), regional variation occurred within the Southeast as well. Proportional losses were similar or less than Southeast and North America losses for 6 BCRs (Table 8). Greater proportional losses occurred in the southernmost BCRs: Peninsular Florida (−60%), and Gulf Coastal Prairie (−29%). Conversely, the largest numeric losses occurred in the Appalachian Mountains (−60 million birds). The species with the greatest numeric losses in Peninsular Florida and the Gulf Coastal Prairie were like those across the Southeast: Red-winged Blackbird, Eastern Meadowlark, and Common Yellowthroat. Common Nighthawk and Northern Mockingbird – both generalists – declined more severely in Peninsular Florida than the Southeast overall, suggesting that common, human-adapted species are not immune to negative anthropogenic habitat effects in the Southeast. Common Nighthawks are impacted by mosquito spraying, as well as the relatively recent conversion of gravel rooftops (i.e., nesting substrate) to smooth, rubber roofs (Brigham et al. 2020), while Northern Mockingbirds are vulnerable to pesticides and other contaminants (Farnsworth et al. 2020). In the Gulf Coastal Prairie, Red-winged Blackbirds and Scissor-tailed Flycatchers (*Tyrannus forficatus*) experienced stronger declines when compared to Southeast-wide estimates of decline. Red-winged Blackbirds are impacted by wetland habitat loss and fragmentation, primarily due to conversion to row crop agriculture, as well as pesticides (Yasukawa and Searcy 2020), while Scissor-tailed Flycatchers are impacted by habitat loss and fragmentation, primarily through loss of shrublands and encroachment and expansion of woodlands (Regosin 2020). Thus, landcover change, particularly expansion of agricultural and loss of shrubby cover, including along fence lines, likely contributed to the sizeable population losses along the northern Gulf Coast.

## CONCLUTIONS

Overall, we found the methodology of Rosenberg et al. (2019) to be readily scalable to the Southeast, and that this analytical approach shows promise as a transparent and repeatable process for evaluating regional avifaunal change. The hierarchical modeling approach yielded robust and defensible estimates of regional population change through time. Further, for the groupings evaluated (e.g., BCR, Habitat, BCC, etc.), model estimates were bounded by reasonably tight confidence intervals (i.e., small margin of error), suggesting the methodology maintained a level of robustness when modeling many species at a regional scale.

It is important to recognize that the methodology employed here is not intended to evaluate population loss or gain at the individual species level. Instead, the hierarchical models generated absolute estimates of loss/gain for evaluation at the group level, across the suite of species assigned to a particular grouping (e.g., BCR, Habitat, Migratory Status, etc.) and category (e.g., migrant vs. resident) within a grouping. Although the methodology fundamentally relies on species-level data, the models draw power (i.e., improved accuracy and precision) by combining abundance estimates from all species assigned to a given group. In so doing, the influence of any single species was dampened except in cases of extremely abundant species (e.g., Blue-gray Gnatcatcher, which was excluded from final analyses) or where a single or few species comprised a given grouping (e.g., Seaside Sparrow was the only species assigned to the Coastal Marsh habitat association). Further, the number of categories within a grouping can also influence the strength of the results. For example, 73% (24 of 33) of the taxonomic families (i.e., categories) had ≤ 5 species. There may be other, less obvious instances where group-level estimates could be strongly influenced by a single or just a few species. Estimates of absolute and proportional gain/loss for any individual species are inherently more sensitive to variability in annual abundance indices (i.e., less robust than group level estimates), especially for species with few observations on BBS routes. Given that the number of species assigned to a group (and by extension the number of categories within a group) influences the power of the analysis, future efforts should carefully weigh tradeoffs between the number of categories and the number of species within each *a-priori* grouping and/or category.

Because our application of hierarchical models required both trend and population estimate for each species within the geography under consideration, our analysis was restricted to species for which published trends and population estimates exist or could be derived for the Southeastern United States (see Methods). This limited our pool of potential species to 146 landbird species (see Methods) representing 1/3rd of the federal trust migratory bird species in the Southeast, thereby highlighting the need to generate robust regional trends and population estimates for other species or taxonomic groups (e.g., shore-birds, colonial waterbirds, secretive marshbirds, etc.). Furthermore, trend estimates used in the assessment of breeding landbirds across the Southeast region are reliant on BBS data, thereby underscoring the importance of maintaining comprehensive coverage of BBS routes within the Southeast Region. For species or seasons (e.g., non-breeding season) where BBS data are not appropriate nor sufficient, support of other national, regional, or broad-scale monitoring programs yielding annual abundance data will be essential (e.g., eBird, regional waterbird atlases). Given the robustness and scalability of the methodology, future assessments could include a wider array of species if and where requisite trend and population estimate data exists.

Importantly, this approach bears promising potential as a transparent and defensible means to track the status of bird populations using a metric – changes in abundance, or degree of population loss or gain – that may be more intuitive and compelling to many stakeholders and decision makers than traditional trend-only results. For example, this approach yields changes in population size to suites of species assigned to *a-priori* habitat groupings thereby elucidating patterns at scales commensurate to those for which management actions and conservation strategies are frequently considered. Magnitude of losses expressed in absolute terms (numbers of birds or percent of populations) may be more evocative and impactful than abstract, trend-based results.

Looking to the future, we recommend periodically repeating these analyses, for example on 5-year cycles, or more frequently based on “moving-window” time periods (e.g., most recent 10 years), and continuing to refine the approach, both conceptually and technically. Outputs from future analyses could help qualify the efficacy of conservation investments and/or expose new patterns or issues worthy of early detection and attention. Further, outputs from such analyses performed on a recurring basis may provide a structured means for: (1) supporting existing priorities and investments; (2) identifying new priorities for investment of staff time and financial resources; and/or (3) evaluating progress towards stated population targets. To that end, we envision the USFWS working in collaboration with other federal, state, and non-governmental organizations to utilize outputs from these assessments to further investigate and understand changes in bird populations to better inform decision making (e.g., investment of resources) as we collectively implement bird-habitat conservation across the Southeast.

### Southeast versus North America

Comparisons of bird population changes between the South-eastern United States and North America yielded similar estimates of population loss for the 141 species evaluated. Analysis by taxonomic family suggested that blackbirds (Icteridae) and new world sparrows (Passerellidae) had the greatest loss in the Southeast, whereas nightjars (Caprimuldidae) and swifts (Apodidae) experienced the greatest proportional losses. These results are consistent with patterns identified by Rosenberg et al. (2019) for North America. Conversely, larks (Alaudidae) and wood warblers (Parulidae) in the Southeast showed less significant population losses compared to estimated losses for these same groups across North America. Further, we observed clear differences in the other group-level analyses between the regional and continental estimates of population change. These regional differences could be influenced by circumstances affecting populations on breeding grounds here in the Southeast, or elsewhere during the annual cycle (i.e., post-breeding, during migration, and/or influenced by issues on the wintering grounds). Understanding drivers behind the observed regional patterns in breeding population loss/gain, and where/why these may differ from patterns continentally, has important implications to conservation investments targeting Southeast breeding bird populations. In that, these population limiting factors may require attention at larger spatial scales or in other regions/ geographies.

### Implications to Conservation Actions in the Southeast

To aid in identifying potential thematic areas warranting further conservation attention across the Southeast, we analyzed suites of birds grouped according to BCR, habitat, taxonomic family, migration status, BCC status, and aerial foraging behavior. Analysis by taxonomic family served to examine population changes within closely related species, as well as changes across the various taxonomic families. Whereas grouping of by foraging behavior and migration status permitted the examination of population change based on life history strategies. Similarly, analysis by habitat associations and BCRs provided a platform to evaluate population change related to the proximate factors (e.g., local and landscape habitat condition, respectively) impacting breeding areas. Further, evaluation of conservation status (BCC) provided a mechanism to assess USFWS priority bird species. While interesting patterns were observed among these groupings, the BCR and habitat groupings likely have the greatest potential in guiding future conservation attention. This is true given that most landbird planning occurs at the scale of a BCR with a focus on specific habitats to guide management recommendations and/or investment decisions.

With that said, we suggest cautious interpretation of our habitat-based results. Because species assignments to specific habitats were not mutually exclusive (i.e., we assigned species to all habitats in which they occurred), the results may be confounded based on the suite of species included within a habitat. A closer examination of species included within a habitat may reveal greater insight to species or groups of species (e.g., Family) that are potentially driving the pattern of population loss or gain within a given habitat. Similarly, additional scrutiny of the species included may reveal known limiting factors that reside outside the Southeast region and facilitate the interpretation of population change within a habitat or other grouping presented here.

Nevertheless, a few habitats and BCRs stood out as warranting immediate attention. From a habitat perspective, birds associated with emergent marsh, grassland/open lands, and shrub-scrub/early successional habitats experienced greater population and proportional losses compared to estimates of loss derived at the North American scale. A closer look at birds associated with shrub-scrub/early seral habitat suggests that early-seral forest habitat species (e.g., Golden-winged Warbler) are experiencing the greatest losses. From a BCR perspective, breeding birds in the Appalachian Mountains, Peninsular Florida and the Gulf Coastal Prairies experienced the greatest declines relative to other BCRs across the Southeast. Interestingly, the Mississippi Alluvial Valley was the only BCR to show relatively stable/potential increase in population size across all species. One potential interpretation of this result, especially considering the trend line since 2010, is that populations may finally be responding positively to intensive and extensive reforestation and other forest habitat conservation activities in the Mississippi Alluvial Valley, which began in the mid-1990s, with the advent of USDA Wetlands Reserve Program (Fig. 9). The lack of precipitous bird population declines (as in most other SE BCRs) and apparent recent increases in overall bird populations in the Mississippi Alluvial Valley coincides with nearly 500,000ac of afforestation reaching >20 years post-planting and an overall gain in forest area exceeding 1,000,000 ac. In other words, it is quite possible that this result is demonstrative of what is possible for bird populations when non-trivial amounts of conservation resources are strategically and consistently focused over a period of several decades. There is no reason to doubt that similar results are possible in other geographies and habitats in the Southeast with focused and dedicated conservation efforts.

**Fig. 9.**
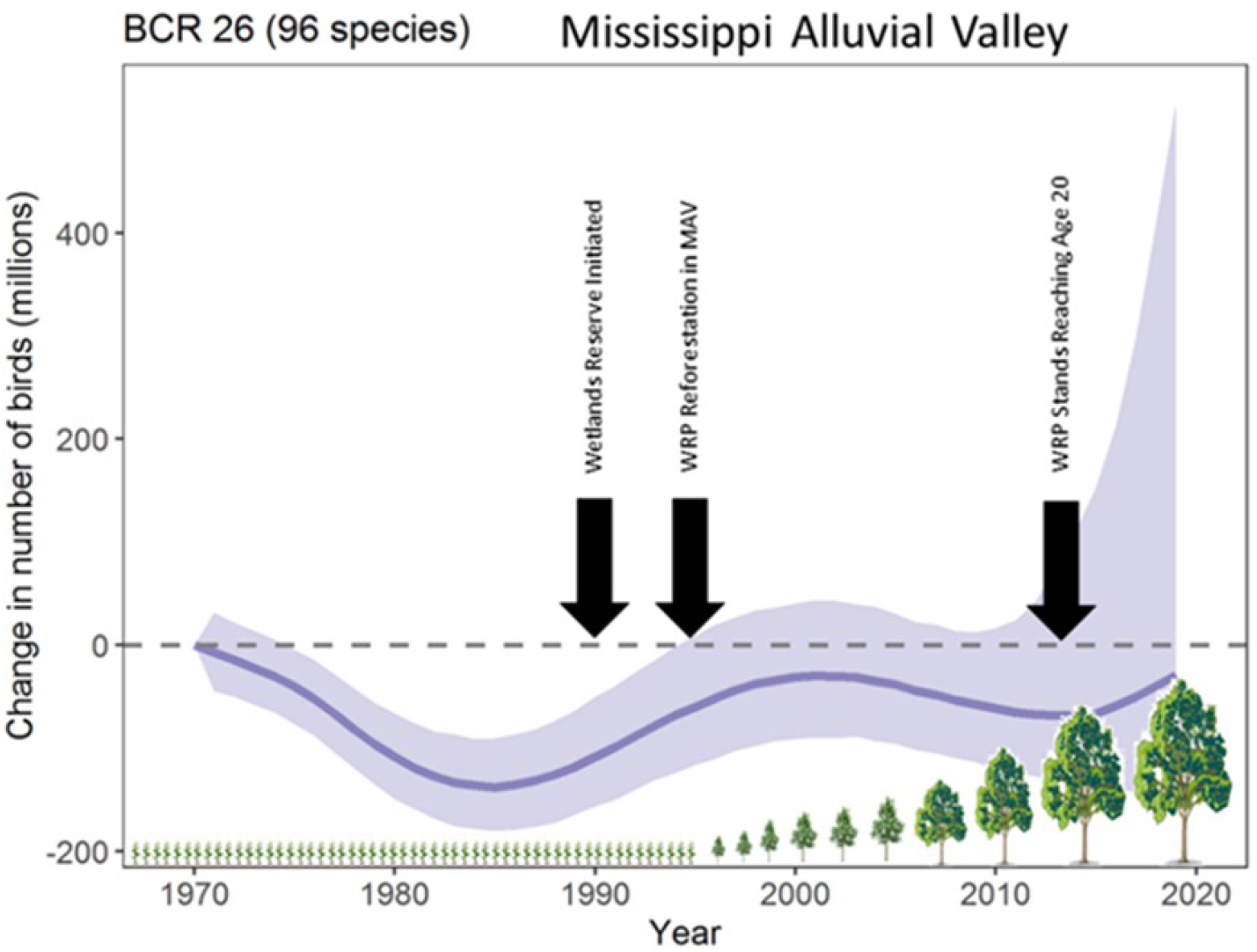
Coincidence of overall bird population change and important conservation actions in the Mississippi Alluvial Valley (BCR 26).

In closing, results suggest that the Rosenberg et al. (2019) methodology of assessing the numerical loss/gain of bird populations is scalable and can be used (with caution) to evaluate population change at finer spatial resolutions and can provide conservation practitioners and decision makers with greater understanding regarding habitats, geographies, or bird groups that may warrant more (or more immediate) attention. The estimated loss of 200 million individuals from Southeast breeding populations since the 1970s is indicative of the alarming magnitude of habitat loss (and habitat fragmentation) and the decline in North America avifauna highlighted by Rosenberg et al. (2019), thus highlighting the urgency for conservation actions.

Results of this assessment reinforced several previously recognized patterns (e.g., significant numerical and proportional decline of grassland birds), as well as to highlight potentially new patterns (e.g., comparatively disproportionate losses of birds in Peninsular Florida) that we had heretofore not recognized. Moving forward, we envision these results providing a basis for further investigating patterns of loss across a range of species x habitat x geographical interactions in a more structured and testable fashion (i.e., patterns of loss/gain as a-priori hypotheses). More specifically, results from these analyses should provide conservation partnerships and initiatives such as Joint Ventures, Eastern Partners in Flight, Northern Bobwhite and Grassland Initiative, and others (e.g., State Wildlife Agencies) with additional information and new perspectives to guide landscape-level planning and on-the-ground bird conservation delivery efforts. In highlighting regional significance of continentally recognized patterns of bird loss, it is our hope that the results presented here provide additional compelling rationale for justifying and bolstering capacity and commitment in curbing losses and “restoring populations” of migratory birds. These results reinforce the urgency to both support and actively participate in in efforts being promoted under the auspices of “*bringing back 3 billion birds*” and the “*Road2Recovery*”, as well as continue to thoughtfully strategize on meaningful ways to effectively address migratory bird declines at scales commensurate with the magnitude of the losses. Engagement and collaboration among broad suites of conservation partners can yield productive outcomes (e.g., as has been the case generally with continental waterfowl populations), but aggregate losses in bird populations suggest that it is cumulatively not enough. Increased capacity, commitment, funding, and collaboration across broad economic and industrial sectors, and society at large will be needed to positively affect land use patterns, and thus, availability of habitat at appropriate spatial scales that allow conservation practitioners to slow, curb, and maybe even reverse widespread declines in bird populations.

