## Supplementary Material - Appendix 2 for "Assessment of Landbird Population Change in the Southeastern United States"

**Table A2.1.** Species grouping criteria, including the level (habitat only), class and definition for each of the six groups: taxonomic family, habitat association (Level I and II), migratory status, conservation status, and aerial insectivores.

| Group | Class | | Definition |
|---|---|---|---|
| **Family** | | | Taxonomic family ($n$ = 36) |
| | See Appendix I | | |
| **Habitat Association** | | | Species primary and/or secondary breeding season habitat association with general (Level I) and finer-scale (Level II) landcover type |
| | Level I | Level II | |
| | Aerial | | Species that exclusively or primarily forage within the air space |
| | Emergent wetland | | Species associated with temporarily or permanently flooded wetlands with standing, rooted, herbaceous plants |
| | | Coastal marsh | Species associated with coastal and estuarine marsh habitat |
| | Forest woodland | | Species associated with forested and woodland habitats |
| | | Eastern deciduous | Species associated with eastern deciduous forest and woodland habitats |
| | | Generalist forest woodland | Species associated with multiple forest and woodland habitats |
| | | Southern pine | Species associated with southern pine forest and woodland habitat |
| | | Spruce-fir / northern hardwood | Species associated with spruce-fir and northern hardwood forest and woodland habitat |
| | Generalist | | Widespread species that use a variety of habitats |

| | | | |
|---|---|---|---|
| | Grassland / open land | | Species associated with habitats dominated by graminoid or herbaceous vegetation, or with limited vegetation |
| | Shrub-scrub / early seral | | Species associated with habitats dominated (i.e., >20% vegetation cover) by shrubs <5 m tall, including true shrubs, young trees in an early successional stage, or trees stunted from environmental conditions |
| | | Early seral forest woodland | Species associated with early seral habitat dominated by tree species in potentially forested areas |
| | | Early seral grassland / open land | Species associated with early seral habitats dominated by graminoid or herbaceous vegetation, or with limited vegetation |
| **Migratory Status** | | | Migratory behavior of species |
| | Migrant | | Long-distance migrant (i.e., breeding and wintering ranges do not overlap) |
| | Partial migrant | | Short-distance migrant (i.e., breeding and wintering ranges distinct but overlapping) |
| | Resident | | Not migratory (i.e., species remains largely within a defined geography year-round) |
| **Conservation Status** | | | Species' Birds of Conservation Concern (BCC 2021) status. Status listed in descending hierarchical order of importance for managers in the southeast from species of greatest concern (Continental) to species of least concern (None) |
| | Continental | | Species listed as a Bird of Conservation Concern at the continental scale |
| | Southeast BCR | | Species listed as a Bird of Conservation Concern for one or more BCRs within the southeast region |
| | Other BCR | | Species listed as a Bird of Conservation Concern for one or more BCRs outside of the southeast region |
| | Not listed | | Species not listed as a Bird of Conservation Concern |
| **Aerial Insectivores** | | | Species that forage for invertebrates within the air space |
| | Exclusive aerial insectivore | | Species that exclusively forage for invertebrates within the air space |
| | Facultative aerial insectivore | | Species that occasionally forage for invertebrates within the air space |
| | None | | Species that rarely or never forage for invertebrates within the air space |

**Table A2.2.** Breeding and non-breeding biomes used as grouping factors in the all-species hierarchical population change model for birds in the southeastern U.S. to incorporate influences across the full annual cycle. Biome names and descriptions from Rosenberg et al. (2019), which were ultimately derived from the Avian Conservation and Assessment Database (ACAD; Partners in Flight 2021).

| Biome | Description |
|---|---|
| **Breeding Biomes** | |
| Aridlands | All arid shrub-dominated communities; primarily in southwestern U.S. and northwestern Mexico; includes ACAD sub-categories of sagebrush, chaparral, desert scrub, barren rocky cliffs, and extensions of tropical dry forest (e.g., thorn-scrub) in southern Texas. |
| Boreal forest | "True" boreal forest of Canada and Alaska; note that some boreal-forest birds also use the boreal zone (primarily spruce-fir) of high mountains in other regions of the U.S. |
| Coasts | All habitats associated with the Coastal zone, including saltmarsh, beach and tidal estuary, mangroves, and rocky cliffs and islands; includes birds that forage primary in the marine zone. |
| Eastern forest | All temperate forest types of eastern U.S. and southeastern Canada (south of the boreal), including northern hardwoods, oak-hickory, pine-oak, southern pine, and bottomland hardwood associations. |
| Forest generalist | Occurs in similar abundance in two or more forest biomes. |
| Grassland | Native grassland, prairie, pasture, and agriculture that supports grassland birds. |
| Habitat generalist | Occurs in similar abundance in three or more major habitat types, usually including forest and non-forest categories. |
| Introduced | Introduced species. |
| Wetland | Freshwater, inland wetlands; does not include coastal marshes or Arctic tundra. |
| **Non-breeding Biomes** | |
| Caribbean | West Indies region, including Cuba, Bahamas, Greater and Lesser Antilles. |
| Coastal | Coastline habitats throughout the western Hemisphere from Arctic to Atlantic and Pacific Coasts of North, Middle, and South America. |
| Introduced | Introduced species. |
| Mexico – Central America | combination of ACAD regions within Mexico and Central America, including *Pacific Lowlands*, *Gulf-Caribbean Lowlands*, *Mexican Highlands*, and species from *Central and South American Highlands* that winter primarily in Central America. |

| | |
|---|---|
| South America | Includes *South American Lowlands*, species from *Central and South American Highlands* that winter primarily in South America, and *Southern Cone* ACAD regions. |
| Southwestern Aridlands | Arid regions of southwestern U.S., northwestern Mexico and Mexican Plateau; included species that winter in arid Chihuahuan grassland habitat. |
| Temperate North America | Broad region encompassing all of Canada and most of the U.S., excluding arid regions in the Southwest. |
| Widespread | Occurs in similar abundance in 3 or more nonbreeding biomes, usually encompassing both temperate North American and Neotropical regions. |
| Widespread Neotropical | Occurs in similar numbers in two or more biome regions within the Neotropics. |
